# Recapitulate Human Cardio-pulmonary Co-development Using Simultaneous Multilineage Differentiation of Pluripotent Stem Cells

**DOI:** 10.1101/2021.03.03.433714

**Authors:** Wai Hoe Ng, Elizabeth K. Johnston, Jun Jie Tan, Jacqueline M. Bliley, Adam W. Feinberg, Donna B. Stolz, Ming Sun, Finn Hawkins, Darrell N. Kotton, Xi Ren

**Author notes:** Correspondence **Authors of correspondence:** Xi Ren, PhD, Carnegie Mellon University, Scott Hall 4N111, 5000 Forbes Avenue, Pittsburgh, PA 15213.

## Abstract

The extensive crosstalk between the developing heart and lung is pivotal for their proper morphogenesis and maturation. However, there remains a lack of model systems for investigating the critical cardio-pulmonary mutual interaction during human embryogenesis. Here, we reported a novel stepwise strategy for directing simultaneous induction of both mesoderm-derived cardiac and endoderm-derived lung epithelial lineages within a single differentiation of human pluripotent stem cells (hPSCs) via temporal specific tuning of WNT and TGF-β signaling in the absence of exogenous growth factors. Using 3D suspension culture, we established concentric cardio-pulmonary micro-Tissues (μTs), and observed expedited alveolar maturation in the presence of cardiac accompany. Upon withdrawal of WNT agonist, the cardiac and pulmonary components within each dual-lineage μT effectively segregated from each other with concurrent initiation of cardiac contraction. We expect our multilineage differentiation model to offer an experimentally tractable system for investigating human cardio-pulmonary interplay and tissue boundary formation during embryogenesis.

## Introduction

Human embryogenesis is a highly orchestrated process that requires delicate coordination between organs originated from different germ layers. Being the two main organs within the chest cavity, the mesoderm-derived heart and endoderm-derived lung have extensive mutual interaction that is essential for their proper morphogenesis.^1–4^ During mouse embryonic development, WNT derived from the second heart field induces specification of pulmonary endoderm, which in turn secretes SHH that signals back to the heart and regulates proper atrial septation.^4–6^ This inter-lineage crosstalk is partly mediated by the multipotent mesodermal progenitors located between the developing heart and lung, which have the potential of lineage contribution to pulmonary endothelium, pulmonary smooth muscle and cardiomyocytes.^1^ However, the translatability of findings derived from rodent models to the understanding of developmental interplay between human cardio-pulmonary systems remains unclear. There is, therefore, a critical need of experimentally tractable systems for investigating human cardio-pulmonary co-development during organogenesis.

Much work has been done for directed differentiation of hPSCs into either cardiomyocytes,^7–13^ or pulmonary epithelium,^14–21^ which often utilizes stepwise differentiation strategies that recapitulate key developmental signaling events. To recapitulate cardiogenesis, hPSCs were sequentially specified into mesoderm, cardiac mesoderm, and then NKX2.5^+^ cardiac progenitors.^7–13^ For pulmonary induction, hPSCs went through stages corresponding to definitive endoderm and anterior foregut endoderm, and then became NKX2.1^+^ lung epithelial progenitors. ^14–21^ Despite significant contribution of these models to the mechanistic understanding of human heart and lung organogenesis, they generally focused on one organ parenchyma at a time. It remains challenging to model and investigate multi-organ co-development within a single differentiation of hPSCs, especially when the organs of interest are derived from different germ layers, as is the case for the heart and lung.

Comparison of existing protocols for single-lineage cardiac and pulmonary differentiation from hPSCs indicates shared regulators despite their distinct germ-layer origin. Firstly, both endodermal and mesodermal specification is facilitated by the inhibition of insulin and phosphoinositide 3-kinase signaling,^7,22,23^ and can be induced by a similar set of paracrine factors, including WNT, BMP and TGF-β.^13,17,24^ It is the quantitative combination of these signaling that determines endoderm versus mesoderm bifurcation.^13,24,25^ This is in consistency with the shared primitive streak origin of both germ layers during gastrulation.^26–28^ Secondly, WNT inhibition not only facilitates the transition from definitive endoderm to anterior foregut endoderm,^24,29^ but also promotes cardiac mesoderm emergence.^7,30–33^ Lastly, retinoic acid (RA) signaling is required for the induction and maturation of both cardiac and pulmonary progenitors.^15,16,23,34,35^ These common paracrine regulation of paralleled cardiac and pulmonary specification is consistent with their close spatial coordinates within the embryonic body planning, as demonstrated by shared HOX genes expression and functional requirement.^36–38^

In this study, we described a stepwise, growth-factor-free protocol for simultaneous induction of cardiac and pulmonary progenitors from a single culture of hPSCs. This is accomplished by initial co-induction of mesoderm and definitive endoderm mixture, followed by their concurrent specification into cardiac (NKX2.5^+^) and lung (NKX2.1^+^) progenitors, respectively, using the same sets of small molecule cocktails modulating WNT and TGF-β signaling in a temporal specific manner. Using 3D suspension culture with continuing WNT activation, we engineered pulmonary-centered, cardio-pulmonary micro-Tissues (μTs), and demonstrated the accompanying cardiac lineage as an essential cellular niche that promoted effective alveolar maturation. Finally, following the withdrawal of WNT agonist, each concentric cardio-pulmonary μT reorganized and ultimately segregated into cardiac-only and pulmonary-only μTs. This work therefore offers an effective hPSC-based model for investigating cardio-pulmonary co-development and tissue segregation during human embryogenesis.

## Results

### Simultaneous induction of cardiac and pulmonary progenitors

Building on existing protocols on cardiac^7–13^ and lung^14–19,39^ differentiation from hPSCs, a stepwise differentiation strategy was developed to enable simultaneous specification of both lineages within a single culture of hPSCs (Fig. 1a). Firstly, a balanced mesodermal and endodermal induction was achieved via fine-tuning of WNT activation in the absence of exogenous supplementation of insulin, TGF-β and BMP4 (Stage-1). Then, a combined inhibition of WNT and TGF-β signaling initiated the specification of the co-induced mesoderm and endoderm towards cardiac and pulmonary specification, respectively (Stage-2). Lastly, reactivation of WNT signaling in the presence of retinoic acid (RA) led to concurrent emergence of NKX2.5^+^ cardiac and NKX2.1^+^ lung progenitors (Stage-3).

**Figure 1:**
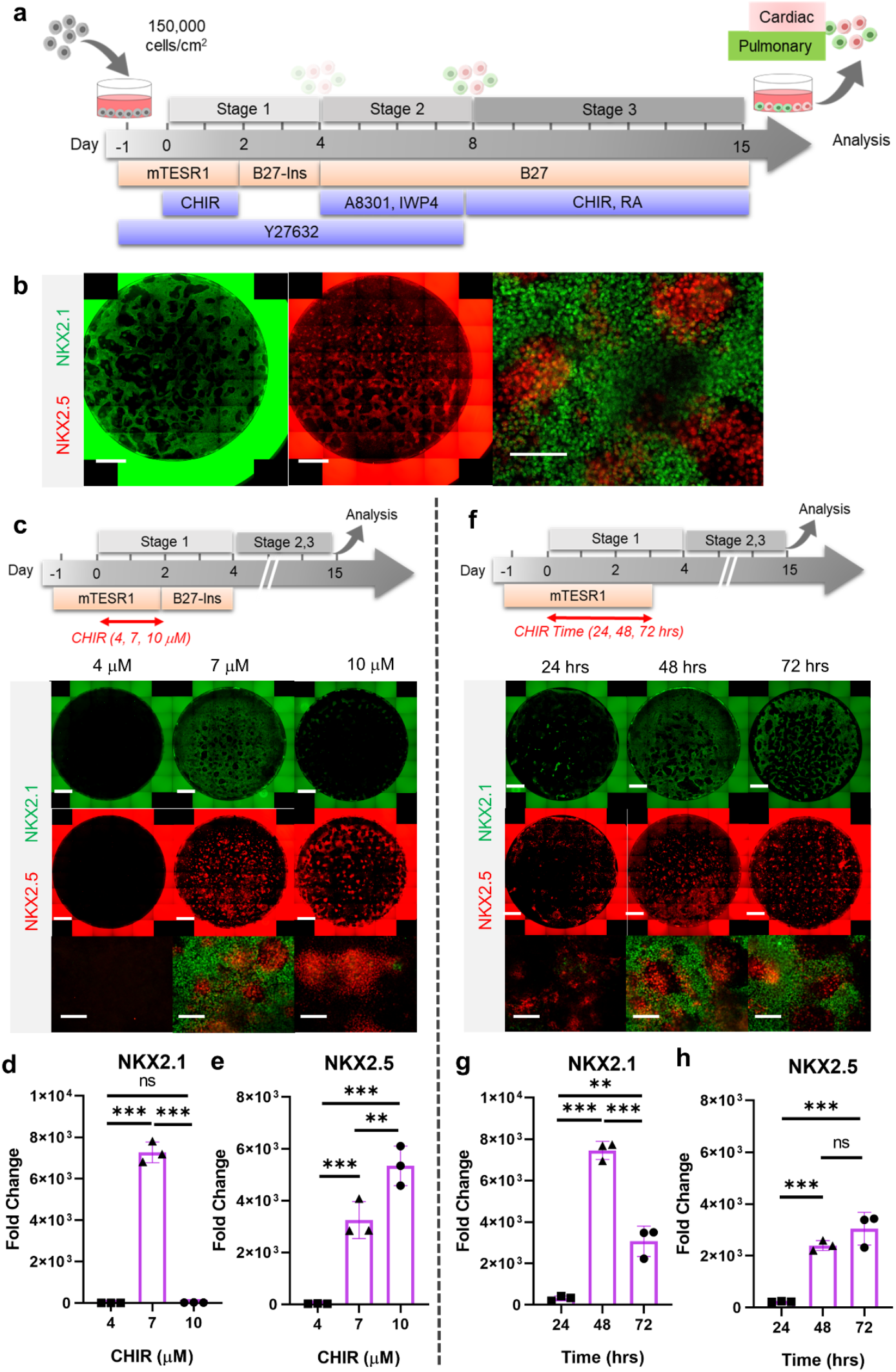
Stepwise cardio-pulmonary co-differentiation from hiPSCs using chemical defined, growth factor-free protocol. (**a**) Schematic diagram showing the overall differentiation strategy. (**b**) Immunofluorescence (IF) showing staining of lung (NKX2.1^+^) and cardiac (NKX2.5^+^). (**c**) IF (**d,e**) and quantitative PCR (qPCR) analysis of the induction of lung and cardiac progenitors on Day-15 of differentiation. (**c-e**) The effects of different CHIR concentrations during Stage-1 of differentiation. Fold change over hiPSCs (**d**) NKX2.1 (n = 3 each; 4 vs. 7, p < 0.001; 7 vs. 10, p < 0.001; 4 vs. 10, p = 0.9993) and (**e**) NKX2.5 (n = 3 each; 4 vs. 7, p < 0.001; 7 vs. 10, p = 0.0053; 4 vs. 10, p = 0.0015). (**f-h**) The effects of different exposure time of CHIR (7 µM) treatment during the first 2 days of differentiation. qPCR analysis of (**g**) NKX2.1 (n = 3 each; 24 vs. 48, p < 0.001; 48 vs. 72, p < 0.001; 24 vs. 72, p < 0.001) and (**h**) NKX2.5 (n = 3 each; 24 vs. 48, p < 0.001; 48 vs. 72, p = 0.1503; 24 vs. 72, p < 0.001). Scale bar = 500 μm for whole well scan; Scale bar = 125 μm for 20X images. All data are mean ± SD. *p < 0.05; **p < 0.01; ***p < 0.001.

WNT signaling is required for mesodermal and endodermal specification in dose-dependent manner during embryogenesis as well as hPSC specification.^7,13,40,41^ Using a human induced pluripotent stem cell (hiPSC) line, BU3 NKX2.1^GFP^; SFTPC^tdTomato^ (BU3-NGST), we examined the possibility of co-inducing mesodermal and definitive endodermal specification solely via the modulation of WNT signaling using a GSK3β inhibitor (CHIR99021, hereafter abbreviated as CHIR) without the addition of exogenous growth factors (e.g. Activin A and BMP4). BU3-NGST were treated with different concentrations of CHIR at 4, 7 and 10 μM for 48 hrs in mTESR1 medium, followed by incubation in growth factor-free differentiation medium (based on RPMI1640 and B-27 minus insulin) (Supplementary Fig. 1a). Towards the end of germ layer induction (Stage-1), we detected co-existence of both definitive endodermal (SOX17^+^) and mesodermal progenitors (MIXL1^+^ and NCAM1^+^), as well as the wide-spread expression of pan-mesendoderm marker (MIXL1) (Supplementary Fig. 1b). In addition, definitive endoderm co-expressed both FOXA2 and SOX17 (Supplementary Fig. 1c). This observation was further confirmed by gene expression analysis of FOXA2, SOX17 and NCAM1 (Supplementary Fig. 1d), which together suggests that 7 μM CHIR drives balanced endoderm and mesoderm induction from hPSCs, while further elevation of CHIR dosage selectively favors mesodermal specification.

To specify the co-induced mesoderm and endoderm towards cardiac and pulmonary lineages, respectively, the Day-4 cells were treated with TGF-β inhibitor (A8301)^35,42,43^ and WNT inhibitor (using IWP4)^7,15^ for 4 days (Stage-2, Day-5 to Day-8), followed by ventralization cocktail consisting of CHIR and RA (essential for lung progenitor specification) for 7 days to Day-15 (Stage 3) (Fig. 1a,b).^35,42,43^ Consistent with CHIR-dependent germ layer induction (Supplementary Fig. 1), the efficiency of cardio-pulmonary specification was tightly regulated by CHIR dosage. We found that on Day-15, cells pre-exposed to CHIR (7 μM) during Stage-1 were able to give rise to robust co-induction of both cardiac (NKX2.5^+^) and pulmonary (NKX2.1^+^) progenitors (Fig. 1c,d,e). In comparison, cells pre-treated with high-CHIR (10 μM) differentiated mainly into the cardiac lineage; while low-CHIR (4 μM) failed to drive effective differentiation of either lineage (Fig. 1c,d,e). We further confirmed the applicability of our cardio-pulmonary co-induction protocol to another independent hiPSC line (BU1) (Supplementary Fig. 2).

The action of CHIR treatment on hPSC differentiation depends on not only its dosage but also the length of exposure time.^44,45^ We evaluated the efficiency of cardio-pulmonary induction following exposure to CHIR (7 µM) for different time spans (24, 48 and 72 hrs), and found that extended CHIR exposure for 48 or 72 hrs was required to induce robust cardio-pulmonary programs (Fig. 1f). Specifically, CHIR favored cardiac specification in a time-dependent manner and plateaued at 48 hrs of treatment (Fig. 1h); while the induction of pulmonary program peaked at 48 hrs of CHIR treatment (Fig. 1g) and declined with further extension of the treatment. Based on these observations, for all subsequent experiments, we used 48-hr treatment of CHIR (7 µM) during Stage-1 of the co-differentiation program. Furthermore, we showed that maintaining hPSCs in mTESR1 Plus during the initial CHIR treatment appeared to be critical for enabling effective cardio-pulmonary differentiation (Supplementary Fig. 3), as compared to using RPMI1640 supplemented with B-27 minus insulin as the basal medium during CHIR treatment.

Exogenous activation of TGF-β and BMP signaling during the very initial steps of hPSC specification has been widely utilized for cardiac^9,13,41^ and pulmonary^15,19,23,42,46^ specification from hPSCs. Here, we investigated how exogenous and endogenous TGF-β and BMP signaling regulates cardio-pulmonary induction during germ layer induction (Stage-1). TGF-β inhibition (using A8301, Day-2 to Day-4) immediately following CHIR treatment abolished both cardiac and pulmonary induction; while TGF-β activation through Activin A supplementation (Day-2 to Day-4) led to pulmonary-only differentiation outcome (Fig. 2a,b,c). This suggests the requirement of endogenous TGF-β for cardio-pulmonary induction and that high-level TGF-β activation favors pulmonary instead of cardiac induction. In parallel, BMP inhibition (using DMH-1) during the same time period compromised cardiac induction and mildly reduced pulmonary specification; while exogenous BMP4 supplementation enhanced cardiac induction but inhibited pulmonary specification (Fig. 2d,e,f). This indicates that endogenous BMP signaling is primarily required for cardiac induction and that exogenous augmentation of BMP signaling further favors the cardiac lineage at the expense of the pulmonary lineage.

**Figure 2:**
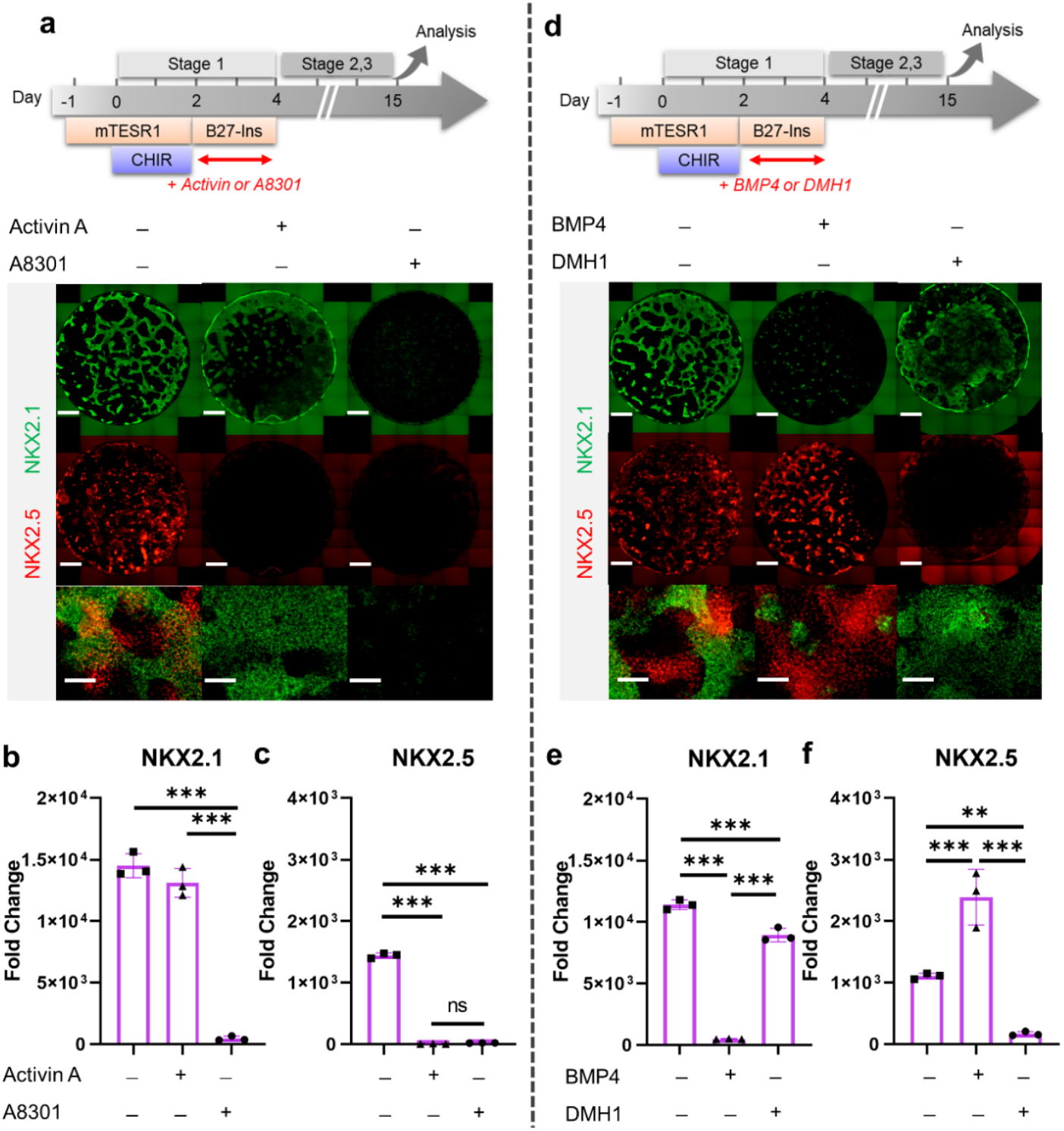
The effect of TGF-β and BMP signaling during Stage-1 of co-differentiation on cardio-pulmonary induction. IF (**a,d**) and qPCR (**b,c,e,f**) analysis of the induction of lung (NKX2.1^+^) and cardiac (NKX2.5^+^) progenitors on Day 15 of differentiation (**a-c**) The effects of exogenous TGF-β activation (Activin A, 20 ng/mL) or its inhibition (A8301, 1 µM). Fold change over hPSCs for (**b**) NKX2.1 (n = 3 each; Activin A^−^ /A8301^−^ vs. Activin A^+^ /A8301^−^, p < 0.001; Activin A^−^ /A8301^−^ vs. Activin A^−^ /A8301^+^, p < 0.001; Activin A^+^ /A8301^−^ vs. Activin A^−^/A8301^+^, p < 0.001) and (**c**) NKX2.5 (n = 3 each; Activin A^−^ /A8301^−^ vs. Activin A^+^ /A8301^−^, p < 0.001; Activin A^−^ /A8301^−^ vs. Activin A^−^ /A8301^+^, p < 0.001; Activin A^+^ /A8301^−^ vs. Activin A^−^ /A8301^+^, p = 0.8649). (**d-f**) The effects of exogenous BMP4 (20 ng/mL) or BMP inhibitor (DMH1, 2 µM). qPCR analysis of (**e**) NKX2.1 (n = 3 each; BMP4^−^ /DMH1^−^ vs. BMP4^+^ /DMH1^−^, p < 0.001; BMP4^−^/DMH1^−^ vs. BMP4^−^ /DMH1^+^, p < 0.001; BMP4^+^ /DMH1^−^ vs. BMP4^−^ /DMH1^+^, p < 0.001) and (**f**) NKX2.5 (n = 3 each; BMP4^−^ /DMH1^−^ vs. BMP4^+^ /DMH1^−^, p < 0.001; BMP4^−^ /DMH1^−^ vs. BMP4^−^ /DMH1^+^, p = 0.0044; BMP4^+^ /DMH1^−^ vs. BMP4^−^ /DMH1^+^, p < 0.001). Scale bar = 500 μm for whole well scan; Scale bar = 125 μm for 20X images. All data are mean ± SD. *p < 0.05; **p < 0.01; ***p < 0.001.

### Shared signaling for cardio-pulmonary co-differentiation from germ-layer progenitors

In previous single-lineage hPSC differentiation studies, TGF-β and WNT inhibition is known to promote pulmonary specification from definitive endoderm ^15,19,35,42,43,46^ as well as the induction of cardiac mesoderm.^7^ Here, we examined how combined inhibition of both TGF-β (using A8301) and WNT (using IWP4) during Day-4 to Day-8 (Supplementary Fig. 4a) regulates cardio-pulmonary specification from germ-layer progenitors established in Stage-1. We found that combined TGF-β and WNT inhibition enhanced both cardiac and pulmonary specification, with TGF-β inhibition having a more profound effect on the cardiac lineage (Supplementary Fig. 4b,c,d). Our finding suggests shared signaling requirement for lung and heart induction from their respective germ-layer progenitors, which is consistent with their close spatial coordinates within the embryonic body planning.^36–38^

In both mouse and human PSC differentiation models, exogenous BMP4 has been shown to be crucial for ventralization of the foregut endoderm to give rise to NKX2.1^+^ lung progenitors. ^15,16,47^ Here in our study, we observed effective cardio-pulmonary co-differentiation in the absence of exogenous BMP4 during ventralization (Stage 3) (Fig. 1b). To address this discrepancy, we investigated how exogenous introduction of BMP4 during ventralization regulated the emergence of cardiac and pulmonary progenitors (Supplementary Fig 5a). Intriguingly, there was no significant differences observed at protein and gene expression level of NKX2.1 and NKX2.5 comparing ventralization in the presence and absence of exogenous BMP4 (Supplementary Fig. 5b,c,d). Nonetheless, endogenous BMP4 was indeed required during this stage of differentiation, as inhibition of BMP4 using DMH1 significantly compromising the induction of both NKX2.1 and NKX2.5 (Supplementary Fig. 5b,c,d).

### 3D suspension culture platform for alveolar induction

To examine whether NKX2.1^+^ lung progenitors derived from the cardio-pulmonary co-induction protocol (Fig. 1a) possess the ability to mature into alveolar type 2 (AT2) epithelial cells, the Day-15 cells were trypsinized and re-plated into the ultra-low adhesion plate for 3D suspension culture (Fig. 3a), and exposed to alveolar maturation medium containing **C**HIR, **K**GF, **D**examethasone, 8-bromoadenosine 3’, 5’-cyclic monophosphate (**c**AMP activator) and **I**BMX (CKDCI).^42,46,48^ Upon transition from 2D to 3D suspension culture in CKDCI medium, the co-induced cardio-pulmonary progenitors self-assembled into pulmonary-centered, concentric, dual-lineage μTs during the overnight culture (Fig. 3b). Following 3 days of 3D suspension culture in CKDCI medium, effective AT2 maturation was observed in the cardio-pulmonary μTs as indicated by robust SFTPC^TdTomato^ fluorescence (Fig. 3c), which sustained up to Day-29 (2 weeks in alveolar maturation, Supplementary Fig. 7a,b). Furthermore, we confirmed lamellar body presence within the induced AT2 cells (Supplementary Fig. 7c). As a control, we cultured Day-15 cardio-pulmonary progenitors on top of the transwell insert for air-liquid interface (ALI) culture or on 2D plastic surface for regular submerged culture, and failed to detect obvious AT2 induction by Day 18 (Fig. 3c Supplementary Fig. 6). Consistent with the observations using fluorescence reporters, NKX2.1 and SFTPC gene expression was significantly upregulated in 3D suspension culture on Day-18 compared to the starting Day-15 cells or cells following ALI maturation (Fig. 3d,e). Our results demonstrated 3D suspension culture as a robust platform to expedite alveolar maturation.

**Figure 3:**
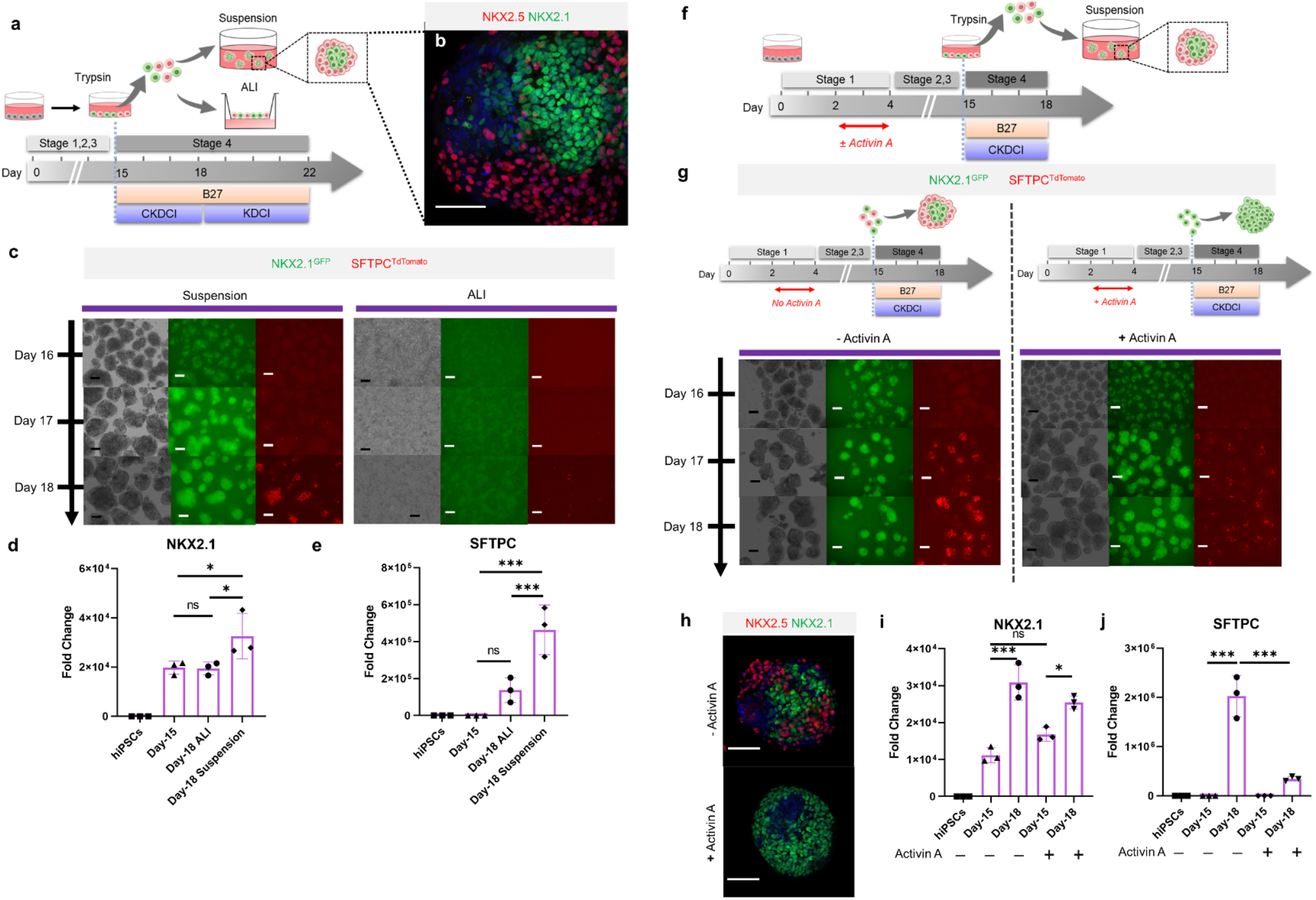
3D suspension culture of cardio-pulmonary μTs expedites AT2 maturation. (**a**) Schematic diagram illustrating the Stage-4 maturation protocol involving replating of Day-15 cardiac and pulmonary progenitors onto ultra-low adhesion plate (for 3D suspension culture or the transwell insert (for ALI culture)). (**b**) Whole mount staining of cardiopulmonary μT on Day-18, scale bar 75 μm. (**c**) Live μT imaging of NKX2.1^GFP^ and SFTPC^TdTomato^ reporter signals during the first 3 days of maturation (Day 16 – Day 18). (**d-e**) qPCR analysis of hiPSC control, Day-15 cells and Day-18 cells (from ALI or suspension culture) for (**d**) NKX2.1 (n = 3 each; Day-15 vs. ALI, p > 0.9999; Day-15 vs. Suspension, p = 0.026; ALI vs. Suspension, p = 0.022), (**e**) SFTPC (n = 3 each; Day-15 vs. ALI, p = 0.9328; Day-15 vs. Suspension, p = 0.0075; ALI vs. Suspension, p = 0.0354) Scale bar = 125 μm. (**f**) Schematic diagram demonstrating experimental design to identify the effect of accompanying cardiac lineage on AT2 maturation. (**g**) Live μT imaging of NKX2.1^GFP^ and SFTPC^TdTomato^ reporter signals during Day 16-18. Scale bar = 125 μm for 10X images. (**h**) Whole mount staining of activin-free and activin-derived μT on Day-18 (**i-j**) qPCR analysis of hiPSC control, Day-15 cells and Day-18 cells (from Activin-free or Activin) for (**i**) NKX2.1 (n = 3 each; Day-15 (No Activin) vs. Day-18 (No Activin), p < 0.001; Day-15 (Activin) vs Day-18 (Activin), p < 0.05; Day-15 (No Activin) vs. Day-15 (Activin), p = 0.1533), (**j**) SFTPC (n = 3 each; Day-15 (No Activin) vs. Day-18 (No Activin), p < 0.001; Day-15 (Activin) vs Day-18 (Activin), p = 0.2417; Day-18 (No Activin) vs. Day-18 (Activin), p < 0.001). All data are mean ± SD. *p < 0.05; **p < 0.01; ***p < 0.001. Scale bar = 125 μm.

To elucidate how the co-induced cardiac lineage modulates the alveolar maturation process, we introduced activin A (20 ng/mL) during germ-layer specification (Fig. 3f), which effectively inhibited mesoderm specification and led to pulmonary-only differentiation outcome on Day-15 (Fig. 2a,b). In the absence of accompanying cardiac cells, although NKX2.1^GFP+^ lung progenitors can be robustly induced and maintained, their alveolar maturation (as indicated by SFTPC^tdTomato^ reporter) following 3 days of maturation in 3D suspension culture was dramatically diminished compared to the cardio-pulmonary group (Fig. 3g). Whole mount imaging of μTs on Day 18 showed pulmonary-only differentiation composed of majority NKX2.1^+^ cells, while cardio-pulmonary μTs were made up of concentric NKX2.1^+^ cells, surrounded by NKX2.5^+^ cells (Fig. 3h). This was further supported by gene expression analysis of NKX2.1 (Fig. 3i) and SFTPC (Fig. 3j). Further extension of CKDCI maturation period for 2 weeks up to Day-29 in the pulmonary-only group failed to produce AT2 induction to a level comparable to the cardio-pulmonary group (Supplementary Fig. 8). This suggests that the cardiac lineage can serve as a cellular niche in supporting alveolar maturation.

### Cardio-pulmonary segregation in the dual-lineage micro-Tissue (μT)

Spatial-temporal regulation of WNT is crucial for early cardiac differentiation,^7,45,49^ however, continuous exposure to WNT activation is known to delay contractile maturation of cardiomyocytes.^50^ In parallel, exogenous WNT activation using CHIR is essential for inducing AT2 maturation and its maintenance until the endogenous AT2 niche is established.^16,51–53^ To investigate how CHIR removal regulates cardio-pulmonary maturation following AT2 establishment on Day-18 in 3D suspension culture (Fig. 4g), we transitioned the maturation medium from CKDCI to KDCI without CHIR (Fig. 4a).^16^ To our surprise, upon CHIR removal, the cardiac and pulmonary components within each dual-lineage μT, which initially arranged in the pulmonary-centered, concentric manner (Fig. 4b), effectively reorganized over time and eventually segregated from each other (Fig. 4a,b).

**Figure 4:**
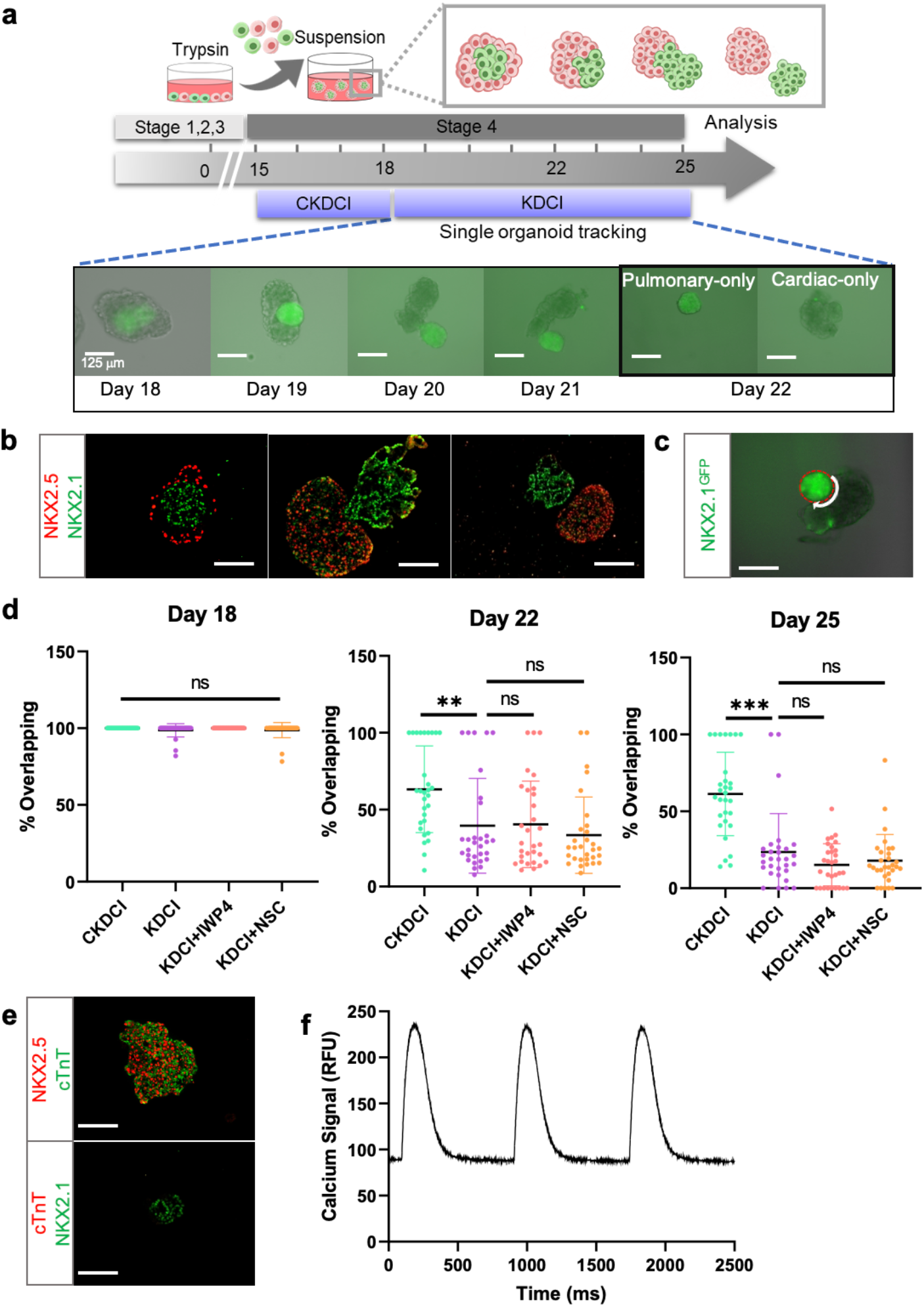
Cardio-pulmonary segregation in the dual-lineage μT. (**a**) Schematic diagram illustrating the timeline for the investigation. (**b**) Histological analysis of cardio-pulmonary μTs at different stages of segregation. (**c**) Diagram showing measurement of the total perimeter of GFP^+^ pulmonary compartment (red color) and its overlapping perimeter with non-GFP compartment (white color) using Image J. (**d**) Scatter plot showing percentage overlapping region of GFP^+^ with non-GFP tissues on Day 18 (n= 30 each;; CKDCI vs. KDCI, p = 0.3979; CKDCI vs. KDCI + IWP4; p > 0.9999; CKDCI vs. KDCI + NSC, p = 0.4293; KDCI vs. KDCI + IWP4, p = 0.3979; KDCI vs. KDCI + NSC, p > 0.9999; KDCI + IWP4 vs. KDCI + NSC, p = 0.4293), Day 22 (n = 30 each; CKDCI vs. KDCI, p = 0.0077; CKDCI vs. KDCI + IWP4; p = 0.0112; CKDCI vs. KDCI + NSC, p < 0.001; KDCI vs. KDCI + IWP4, p = 0.9994; KDCI vs. KDCI + NSC, p = 0.8318; KDCI + IWP4 vs. KDCI + NSC, p = 0.7674) and Day 25 (n = 30 each; CKDCI vs. KDCI, p < 0.001; CKDCI vs. KDCI + IWP4; p < 0.001; CKDCI vs. KDCI + NSC, p < 0.001; KDCI vs. KDCI + IWP4, p = 0.4271; KDCI vs. KDCI + NSC, p = 0.7275; KDCI + IWP4 vs. KDCI + NSC, p = 0.9623). (**e**) Histological analysis of cTnT expression on the segregated cardiac and pulmonary μTs, with co-staining of NKX2.5 and NKX2.1. (**f**) Calcium influx measured using calcium indicator, Xrhod-AM. Scale bar = 125 μm for 20X images. All data are mean ± SD. *p < 0.05; **p < 0.01; ***p < 0.001.

To quantitatively assess this segregation process, we performed time-lapse single-μT tracking and calculated the percentage of overlapping between the cardiac and pulmonary tissues by measuring the length of overlapping border between the GFP^+^ and non-GFP components and normalizing it by the total perimeter of the GFP^+^ pulmonary component (Fig. 4c). We compared the segregation process in the presence (CKDCI) and absence (KDCI) of CHIR, and found that although cardio-pulmonary segregation took place in both medium conditions, it was significantly expedited by the withdrawal of CHIR (Fig. 4d). To investigate the requirement of endogenous WNT signaling for this segregation process, we introduced inhibitors of canonical (IWP4) and non-canonical (NSC668036, a Dishevelled inhibitor) WNT signaling, ^54^ and did not detect obvious difference in the segregation process as compared to the control KDCI condition (Fig. 4d). In parallel with the cardio-pulmonary segregation, cardiac contraction was observed 7 days following CHIR withdrawal (Supplementary Video 1). Immunohistochemical analysis demonstrated specific co-expression of NKX2.5 and cardiac troponin T (cTnT) in the segregated cardiac μT (Fig. 4e). The contractile function of the segregated cardiac μT was further confirmed via the detection of calcium influx (Fig. 4f, Supplementary Video 2).

## Discussion

Here, we described a novel strategy to model human cardio-pulmonary co-development using multi-lineage hPSC differentiation. We demonstrated that upon co-induction of mesoderm and endoderm, a series of shared signaling events were capable of driving simultaneous cardiac and pulmonary specification from their respective germ-layer progenitors. Transitioning the co-induced cardiac and pulmonary progenitors to 3D suspension culture, we observed expedited alveolar maturation within 3 days, which was supported by the accompanying cardiac lineage. In 3D suspension culture, each cardio-pulmonary μT effectively segregates into separate cardiac and pulmonary μTs, which was partially inhibited by WNT activation. This study therefore delivers an effective *in vitro* model for studying the mechanistic interplay between developing heart and lung during human embryogenesis.

The extensive cardio-pulmonary mutual interaction during organogenesis has been well documented in the mouse model,^1,2,4^ however, the translatability of these findings to human embryogenesis remains elusive due to the lack of proper model systems. Human PSC differentiation has offered an effective means for recapitulating and investigating human organogenesis, and tremendous progresses have been made towards directed cardiac or pulmonary specification.^7–23,39^ However, almost all existing models have been focusing on one parenchymal lineage at a time, and therefore lack the ability to support investigation on inter-organ crosstalk. Here, building on the established understanding of signaling events necessary for cardiac and pulmonary induction,^7–23,39^ we have developed a robust protocol for simultaneous cardio-pulmonary co-differentiation from hPSCs. Within our co-differentiation system, unrestricted interaction between cells of both lineages is enabled even before their lineage commitment.

Most current attempts on pulmonary induction from hPSCs relies on initial TGF-β activation using growth factor (Activin A) supplementation, which is critical for definitive endoderm specification. Here, we showed that by fine tuning of WNT signaling using a small-molecule inhibitor of GSK-3β (CHIR), robust induction of endoderm and subsequently lung progenitors can be achieved without any exogenous growth factor. This is consistent with the observation that CHIR was capable of inducing cardiac differentiation in replacement of combined effect of exogenous Activin A and BMP4.^7^ Nonetheless, Nodal and BMP signaling remains crucial in mesoderm and endoderm specification, as inhibition of these signaling abolished effective cardio-pulmonary co-induction.

Our study demonstrated the requirement of endogenous TGF-β signaling for effective cardio-pulmonary induction, as well as the critical role of endogenous BMP signaling for cardiogenesis. Furthermore, we found that temporal-specific action of the same set of small molecules regulating TGF-β and WNT signaling was capable of driving mesoderm-to-cardiac and endoderm-to-pulmonary specification in a concurrent manner. Moreover, BMP4 has been shown to improve NKX2.1^+^ lung progenitor specification in both mouse and human iPSCs.^15,16,47^ In our system, endogenous instead of exogenous BMP signaling was required during a developmental stage corresponding to foregut ventralization for effective co-emergence of cardiac and pulmonary progenitors. This is in line with the close spatial positioning of developing heart and lung primordia within embryonic body patterning, which implies their exposure to a similar paracrine microenvironment.^4,55^

To achieve alveolarization, NKX2.1^+^ lung progenitors are usually embedded in extracellular matrices, such as Matrigel and collagen.^42,46,48^ Here, we established an effective approach that enabled AT2 cell maturation within 3 days in suspension culture of 3D cell aggregates spontaneously formed from Day-15 cardiac and pulmonary progenitors. We further demonstrated that the presence of accompanying cardiac lineage is critical for robust alveolar induction. This observation is consistent with the recently reported inter-dependence between cardiac and pulmonary lineages during embryogenesis.^4^ In addition, the presence of mesoderm-derived stromal cells has been shown to be essential for effective alveolarization *in vivo*^55–57^ and *in vitro*.^19,43^ Furthermore, cells of the mesodermal lineage are known to be robust producers of extracellular matrix, which may also contribute to the effective alveolar maturation in the absence of external extracellular matrix support. The ability to enable effective alveolar induction from hiPSC-derived lung progenitors in a convenient suspension culture also opens the door to large scale production of alveolar cells, a critical enabling step for regenerative medicine applications.

Using dual-lineage cardio-pulmonary μTs formed from the co-induced progenitors, we observed a novel process of cardio-pulmonary tissue segregation. The human body cavities are highly crowded spaces, filled with different tissues and organs that are in close contact with each other. It remains enigmatic how inter-organ boundaries are maintained to prevent undesired cell migration or tissue merging. Our cardio-pulmonary tissue segregation model suggests an intrinsic mechanism that effectively establishes a boundary between two distinct parenchymal lineages even when they are initially mingled together. Although no model of collective migration has been described in the context of cardio-pulmonary development, studies in other model systems suggest cell-cell communication and paracrine signaling (e.g. WNT) to be crucial for directed cell migration during development.^58–60^ Here we found that exogenous WNT activation via GSK-3β inhibition effectively slowed down the cardio-pulmonary segregation, while inhibition of endogenous WNT (canonical and non-canonical) did not obviously affect the process. In consistence with our observation, it has been shown that inhibition of non-canonical WNT signaling does not stop collective cell migration but distorting migration direction.^54^

In conclusion, our work offers a novel model for investigating the molecular and cellular mechanisms underlying human cardio-pulmonary co-development and tissue boundary formation. We also expect this work to be of potential use for studying congenital diseases affecting both cardiovascular and pulmonary systems, such as congenital diaphragmatic hernia.

## Materials and Methods

### Materials

Detailed information regarding reagents for culture and differentiation medium was summarized in Supplementary Table 1. Reagents, equipment, and probes for quantitative PCR (qPCR) analysis were summarized in Supplementary Table 3. Antibodies and Reagents for immunofluorescence staining were summarized in Supplementary Table 4.

### Maintenance of human induced pluripotent stem cells (hiPSCs)

The BU3-NGST hiPSC line was obtained as a kind gift from the laboratories of Dr. Darrell Kotton and Dr. Finn Hawkins (Boston University). BU3 hiPSC line was derived from a healthy donor and carries both NKX2.1^GFP^ (NG) and Surfactant protein C (SFTPC)^tdTomato^ (ST) reporters.^42,43^ hiPSCs were maintained on Matrigel-coated (ESC-qualified) 6-well tissue culture plate with mTESR1 Plus medium with regular medium changed every other day. hiPSCs passaging was performed every 5-7 days using ReLESR at a plating ratio of 1:10. All cells used in this study were tested negative for mycoplasma contamination using Universal Mycoplasma Detection Kit (ATCC, 30-1012K).

### Simultaneous induction of cardiac and pulmonary progenitors from hPSCs

hiPSCs maintained in mTESR Plus were dissociated into single cells using StemPro Accutase. 150,000 cells/cm^2^ on hESC-qualified Matrigel-coated 96-well plate, and cultured in mTESR Plus supplemented with 10 μM Y-27632 (ROCK inhibitor) for 24 hrs prior to differentiation. The overall protocol for stepwise cardio-pulmonary co-differentiation was summarized in Supplementary Table 2. To induce a balanced mixture of mesodermal and definitive endodermal cells, hiPSCs were first incubated in mTESR Plus medium supplemented with different concentration (4, 7, 10 μM) of CHIR99021 (GSK3β inhibitor) and 10 μM Y-27632 for 48 hrs. This was followed by an additional 48 hrs incubation in serum-free differentiation medium consisting of RPMI 1640 supplemented with 2% B-27 minus insulin, 1 x GlutaMAX and 10 μM Y-27632. In some experiments, Activin A (20 ng/mL), BMP4 (20 ng/mL), A8301 (TGF-β inhibitor, 1 μM) or DMH-1 (BMP inhibitor, 2 μM) were introduced to examine how TGF-β and BMP signaling regulated mesodermal and endodermal specification. Differentiation outcomes were assessed by immunostaining and qPCR analysis of mesodermal (NCAM1) and definitive endodermal (SOX17) markers.

Following Stage-1, all subsequent differentiation procedures were performed using medium recipes formulated based on RPMI 1640 medium supplemented with 2% B-27 and 1x GlutaMAX, referred to as ‘basal medium’. To initiate simultaneous cardiac and pulmonary specification, Day-4 cells were incubated for 4 days in Stage-2 medium, containing basal medium supplemented with 1 μM A8301, 5 μM IWP4 and 10 μM Y-27632. In some experiments, co-differentiation medium without either A8301 or IWP4 was utilized to investigate the impact of inhibition of TGF-β and WNT signaling.

Following Stage-2, to induce simultaneous specification of both cardiac and lung progenitors, co-differentiating cells were incubated for 7 days in Stage-3 medium containing basal medium supplemented with 3 μM CHIR99021 and 100 nM Retinoic acid (RA). Green fluorescence of the NKX2.1^GFP^ reporter was examined daily using EVOS FL Auto 2 Imaging System to monitor the emergence of lung progenitors. On Day-15 of co-differentiation, the expression of cardiac (NKX2.5) and lung (NKX2.1) progenitor markers was evaluated by immunofluorescence staining and qPCR.

### Co-maturation of cardio-pulmonary progenitors in air-liquid interface (ALI) culture

On Day-15 of cardio-pulmonary co-differentiation, cells were dissociated into single cells using TrypLE Express, and re-plated at 500,000 cells/cm^2^ onto the apical side of each 24-well Transwell insert (pore size of 0.4 μm, pre-coated with 1% growth factor-reduced Matrigel) in 100 μL maturation medium. Basolateral side of the transwell insert was filled with 500 μL of maturation medium. The maturation medium was basal medium supplemented with 3 μM **C**HIR99021, 10 ng/mL Keratinocyte growth factor (**K**GF), 50 nM **D**examethasone, 0.1 mM 8-bromoadenosine 3’, 5’-cyclic monophosphate (**c**AMP, AMP-activated protein kinase activator) and 0.1 mM 3-**i**sobutyl-1-methylxanthine (IBMX, PKA activator), which was referred to as **CKDCI** medium. 10 μM Y-27632 was added during the initial 24 hrs following re-plating. The next day, all medium on the apical side was removed. 200 μL of fresh CKDCI medium without Y-27632 was added to the basolateral side to establish ALI culture, and was replaced daily. Red fluorescence from the SFTPC^TdTomato^ reporter was examined daily using EVOS Imaging System to monitor the emergence of alveolar type 2 (AT2) cells. On Day-3 of ALI maturation, Transwell membrane were excised from the insert, and analyzed by qPCR (NKX2.1, SFTPC).

### Co-maturation of cardio-pulmonary μTs in 3D suspension culture

On Day-15 of cardio-pulmonary co-differentiation, cells were dissociated into single cells using TrypLE Express. A total of 250,000 cells in 500 μL CKDCI maturation medium was transferred into each well of 24-well ultra-low adherence plate and cultured with agitation at 125 rpm to form cardio-pulmonary μTs. 10 μM Y-27632 was added during the initial 24 hrs following re-plating. Following 3 days of culture in CKDCI medium, CHIR99021 was removed and μT culture was continued in KDCI medium for an additional 7 days. At desired time points of 3D suspension maturation, μTs were analyzed by histology (NKX2.1, NKX2.5, cTnT) and qPCR analysis (NKX2.1, SFTPC).

### Single μT time-lapse imaging and analysis

To investigate the segregation of cardio-pulmonary μTs into their respective cardiac and lung μTs, following 3 days suspension culture in CKDCI medium in 24-well ultra-low adherence plate, single μT was transferred into each well in 96-well ultra-low adherence plate and cultured for an additional 7 days. The following medium recipes were examined for cardio-pulmonary segregation: KDCI medium, KDCI medium supplemented with 3μM CHIR99021, KDCI medium with 5 μM IWP4, and KDCI medium with 50 μM NSC668036. Time-lapse imaging was performed on Day-18, Day-22 and Day-25 following μT transfer to monitor the segregation process. The pulmonary compartment within each cardio-pulmonary μT was tracked based on the NKX2.1^GFP^ reporter. To quantify the segregation between the two compartments within each μT. Image J was used to measure the overlapping perimeter between GFP^+^ (pulmonary) and non-GFP (cardiac) compartments, which was then normalized to total perimeter of GFP^+^ compartments and expressed as the percentage of overlapping.

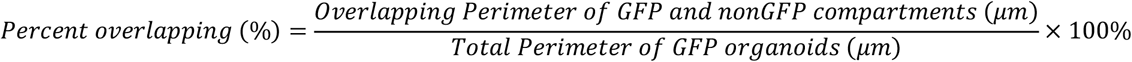

### qPCR analysis

Total RNA was extracted using TRIzol, processed by chloroform extraction, precipitated using 1 volume of absolute isopropanol with 50 μg/mL of RNase-free glycoblue as carrier, washed with 75% ethanol, air-dried, solubilized in RNase-free water and quantified using NanoDrop 2000 spectrophotometer. cDNA was synthesized via reverse transcription of 1 μg total RNA with random hexamers and the High-Capacity cDNA Reverse Transcription kit according to manufacturer’s instruction. Real-time qPCR analysis was performed on CFX96 Touch Real-Time PCR Detection System using TaqMan probes. Each reaction mixture was prepared by combining 1 μL of probe, 10 μL of TaqMan Master Mix, 1 μL of cDNA (equivalent to 50 ng), and the final volume was brought up to 20 μL. The final Ct value was normalized to housekeeping gene (β-actin), using comparative Ct method. Unless otherwise specified, baseline, defined as fold change = 1, was set as undifferentiated hiPSCs, or if undetected, a cycle number of 40 was assigned to allow fold change calculations.^42^ List of TaqMan probes was summarized in Supplementary Table 3.

### Immunofluorescence staining on 2D cell samples

Cells were fixed with ice-cold methanol, air-dried, rehydrated with phosphate-buffered saline (PBS), permeabilized with 1% (v/v) Triton X-100, blocked in 1% (w/v) bovine serum albumin in PBS (blocking buffer), incubated with primary antibodies diluted in blocking buffer at 4°C overnight, and incubated with corresponding fluorescence-conjugated secondary antibodies in blocking buffer at room temperature (RT) for 45 min. Nuclear counterstain was performed using Hoechst-33342 (1:500) in PBS. Fluorescence images were acquired using EVOS Imaging System. All antibodies used and their respective dilution were summarized in Supplementary Table 4.

### Histology

The μTs were fixed with 4% paraformaldehyde, embedded in HistoGel and then in paraffin. Tissue processing and paraffin embedding was performed in Research Histology Lab of Pitt Biospecimen Core at the University of Pittsburgh Medical Center (UPMC) Shadyside Hospital. Paraffin blocks were sectioned at 5 μm thickness, transferred onto glass slides, rehydrated by sequential incubation in Histoclear, 100% ethanol, 95% ethanol and distilled water. To unmask antigen, slides were treated with Antigen Unmasking Solution at 95°C for 20 min and cooled down to RT. Immunofluorescence staining was performed as described above for 2D cell samples. After the final wash, slides were mounted with DAPI Fluoromount-G, and imaged using EVOS Imaging System. All antibodies used and their respective dilution were summarized in Supplementary Table 4.

### Contraction and calcium signal

To assess contraction of cardiac μT, segregated cardiac μT was stained with 5 μM of Cal-520 AM (AAT Bioquest, 21130), a calcium indicator dye. Calcium imaging (500 frames per second) was performed using a Prime 95B Scientific CMOS camera (Photometrics) mounted on an epifluorescent stereomicroscope (Nikon SMZ1000) with a GFP filter and an X-cite Lamp (Excelitas).

### TEM

Cardio-pulmonary μTs were fixed in 2.5% glutaraldehyde in 0.1 M PBS (pH7.4) for at least 1 hr. After 3 washes in 0.1 M PBS for 10 min each, the μTs were post fixed in 1% Osmium tetroxide containing 1% potassium ferricyanide at 4°C for 1 hr, followed by 3 washes in 0.1 M PBS for 10 min each. μTs were dehydrated in graded series of ethanol starting from 30%, 50%, 70%, 90% and finally 100% of ethanol for 10 min each. μTs were further dehydrated epon for 1 hr at RT. This step was repeated for another three times prior to embedding in pure epon at 37°C for 24 hrs. Finally, the μTs were cured for 48 hrs at 60°C. The presence of lamellar body in cardio-pulmonary μTs were identified using JEM 1400 Flash TEM.

### Statistics

Statistical methods relevant to each figure were outlined in the accompanying figure legend. At least three biological replicates were performed for each group under comparison. Unless otherwise indicated, unpaired, 2-tailed Student’s t tests were applied to comparisons between two groups. For comparisons among three or more groups, one-way ANOVA was performed followed by Tukey multiple comparison tests. Results are displayed as mean ± SD, with p < 0.05 considered statically significant. n values referred to biologically independent replicates.

## Data availability

The authors declare that all data supporting the findings of this study are available within the article and its supplementary material files, or from the corresponding author on reasonable request.

## Acknowledgments

This work was supported by Samuel & Emma Winters Foundation A025662 (to X.R.) and the Department of Biomedical Engineering and College of Engineering at Carnegie Mellon University. We are grateful to Drs. Yu-li Wang and David Li for advice on collective cell migration, to Misti West for laboratory management, and to Anthony Green and the Pitt Biospecimen Core at the University of Pittsburgh for assistance with histology.

## Author’s contributions

X.R. and W.H.N. designed the project and wrote the manuscript. W.H.N, E.K.J and J.M.B performed the experiments and analyzed data. W.H.N., X.R., J.J.T., J.M.B., A.W.F. and F.H. interpreted data. D.B.S. and M.S. performed the electron microscopy analysis. F.H. and D.N.K. provided the BU3-NGST and BU1 hiPSC lines and advised on pulmonary differentiation. J.J.T., E.K.J., J.M.B. and F.H. edited the manuscript.

## SUPPLEMENTARY INFORMATION

**Supplementary Table 1:**
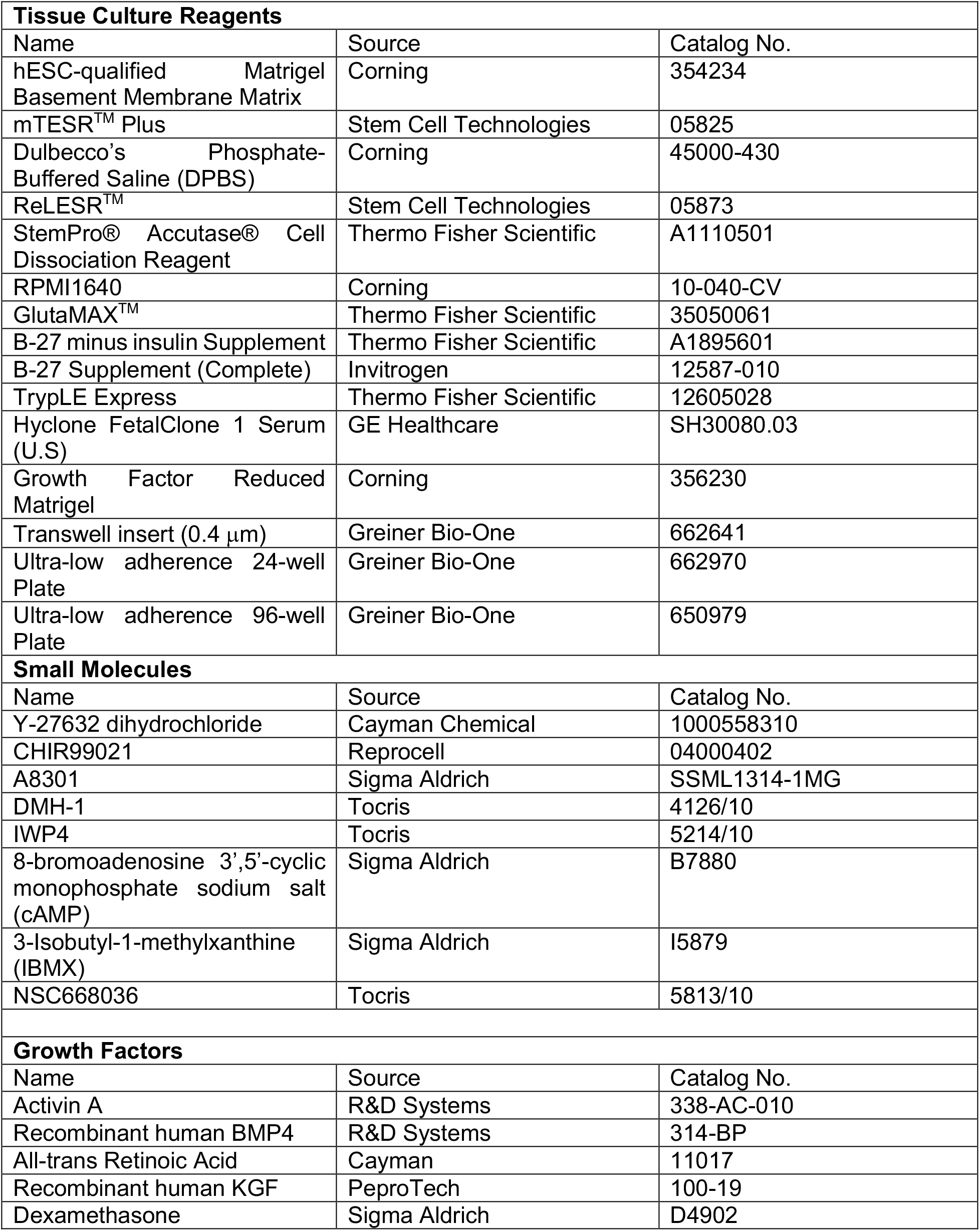
Tissue culture reagents, small molecules and growth factors.

**Supplementary Table 2:** Media Recipes/Composition.

| Media | Base | Cytokines/Growth Factors | Final Concentration |
| --- | --- | --- | --- |
| Stage 1: Day 0 - 1 | mTESR Plus | CHIR99021 | 7 μM |
| | | Y27632 | 10 μM |
| Stage 1: Day 2 – 3 | RPMI 1640 | Y27632 | 10 μM |
|  | B-27 minus insulin |  |  |
|  | GlutaMAX (1x) |  |  |
| Stage 2: Day 4 – 7 | RPMI 1640 | A8301 | 1 μM |
| | B-27 complete | IWP4 | 5 μM |
| | GlutaMAX (1x) | Y27632 | 10 μM |
| Stage 3: Day 8 – 14 | RPMI 1640 | CHIR99021 | 3 μM |
|  | B27 complete | Retinoic acid | 100 nM |
|  | GlutaMAX (1x) |  |  |
| Stage 4: Day 15 -17 | RPMI 1640 | CHIR99021 | 3 μM |
|  | B27 complete | KGF | 10 ng/mL |
|  | GlutaMAX (1x) | Dexamethasone | 50 nM |
|  |  | cAMP | 0.1 mM |
|  |  | IBMX | 0.1 mM |
| Stage 4: Day 18 onwards | RPMI 1640 | KGF | 10 ng/mL |
|  | B27 complete | Dexamethasone | 50 nM |
|  | GlutaMAX (1x) | cAMP | 0.1 mM |
|  |  | IBMX | 0.1 mM |

**Supplementary Table 3:**
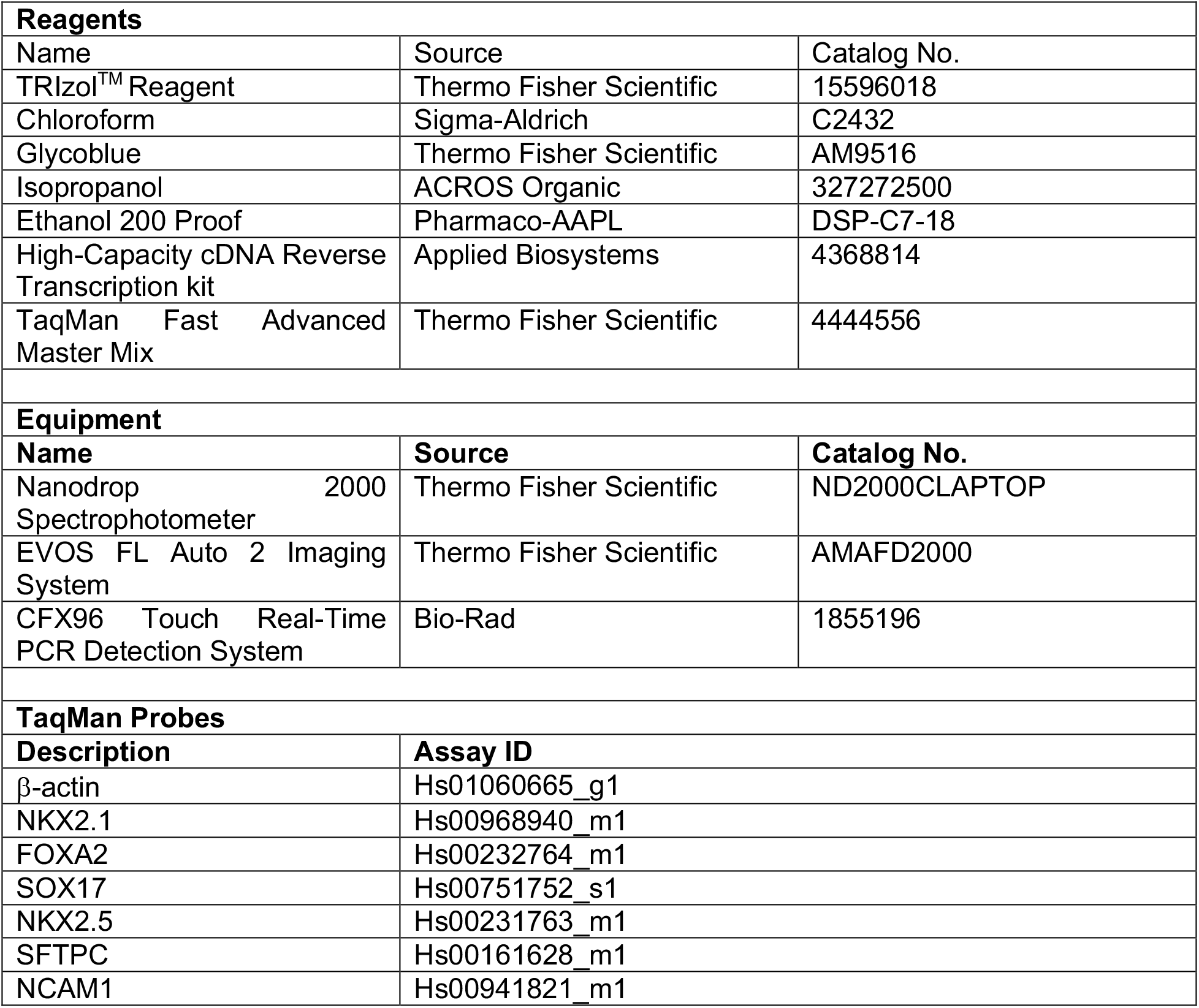
Reagents, equipment and probes for qPCR.

**Supplementary Table 4:**
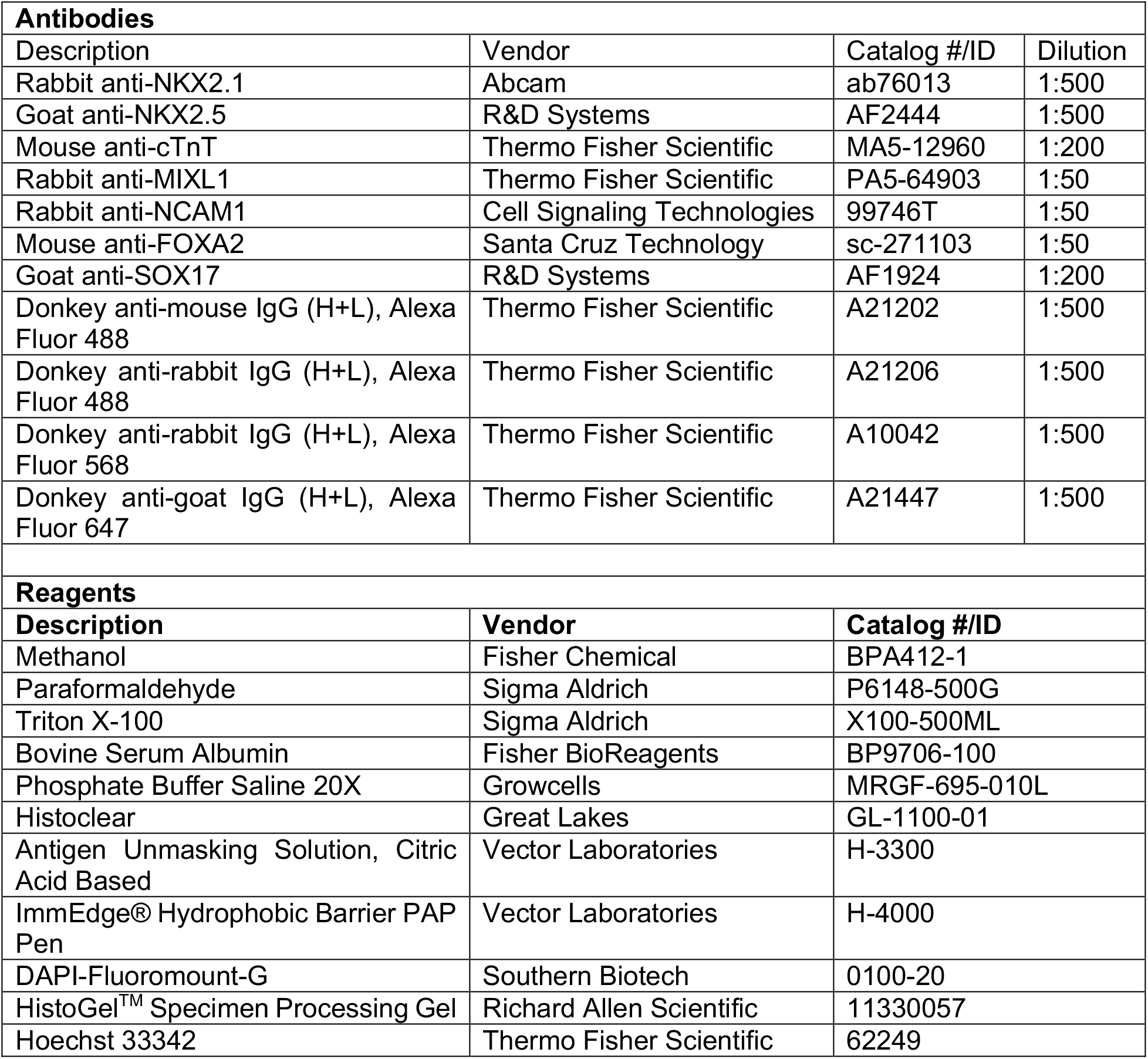
Antibodies and Reagents for Immunostaining.

| Antibodies |  |  |  |
| --- | --- | --- | --- |
| Description | Vendor | Catalog #/ID | Dilution |
| Rabbit anti-NKX2.1 | Abcam | ab76013 | 1:500 |
| Goat anti-NKX2.5 | R&D Systems | AF2444 | 1:500 |
| Mouse anti-cTnT | Thermo Fisher Scientific | MA5-12960 | 1:200 |
| Rabbit anti-MIXL1 | Thermo Fisher Scientific | PA5-64903 | 1:50 |
| Rabbit anti-NCAM1 | Cell Signaling Technologies | 99746T | 1:50 |
| Mouse anti-FOXA2 | Santa Cruz Technology | sc-271103 | 1:50 |
| Goat anti-SOX17 | R&D Systems | AF1924 | 1:200 |
| Donkey anti-mouse IgG (H+L), Alexa Fluor 488 | Thermo Fisher Scientific | A21202 | 1:500 |
| Donkey anti-rabbit IgG (H+L), Alexa Fluor 488 | Thermo Fisher Scientific | A21206 | 1:500 |
| Donkey anti-rabbit IgG (H+L), Alexa Fluor 568 | Thermo Fisher Scientific | A10042 | 1:500 |
| Donkey anti-goat IgG (H+L), Alexa Fluor 647 | Thermo Fisher Scientific | A21447 | 1:500 |
| Reagents |  |  |  |
| Description | Vendor | Catalog #/ID |  |
| Methanol | Fisher Chemical | BPA412-1 |  |
| Paraformaldehyde | Sigma Aldrich | P6148-500G |  |
| Triton X-100 | Sigma Aldrich | X100-500ML |  |
| Bovine Serum Albumin | Fisher BioReagents | BP9706-100 |  |
| Phosphate Buffer Saline 20X | Growcells | MRGF-695-010L |  |
| Histoclear | Great Lakes | GL-1100-01 |  |
| Antigen Unmasking Solution, Citric Acid Based | Vector Laboratories | H-3300 |  |
| ImmEdge® Hydrophobic Barrier PAP Pen | Vector Laboratories | H-4000 |  |
| DAPI-Fluoromount-G | Southern Biotech | 0100-20 |  |
| HistoGel™ Specimen Processing Gel | Richard Allen Scientific | 11330057 |  |
| Hoechst 33342 | Thermo Fisher Scientific | 62249 |  |

**Supplementary Fig. 1:**
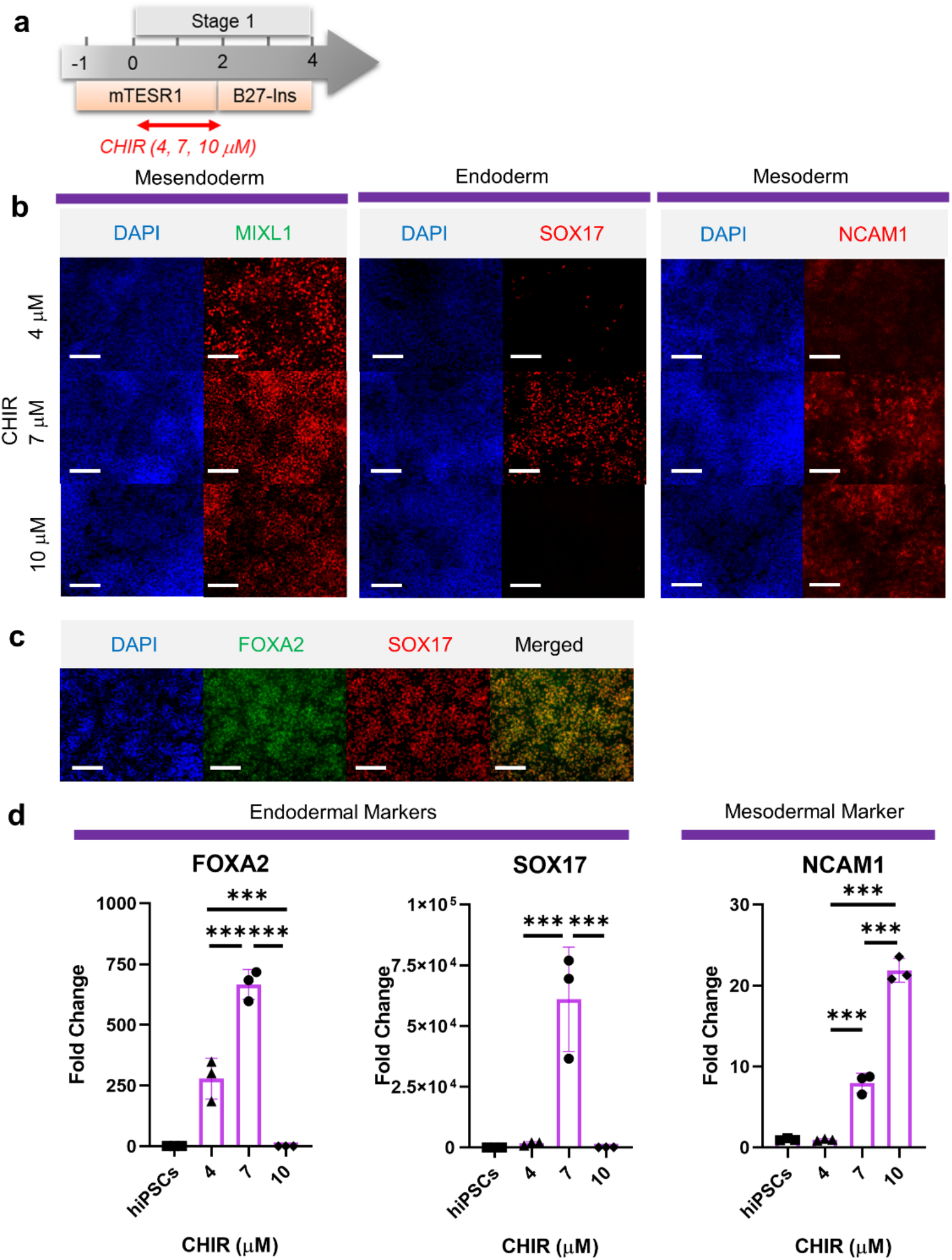
Mesoderm and endoderm co-induction from hPSCs using CHIR. (**a**) Diagram showing the experimental design. (**b**) Cells following Stage-1 differentiation expressed MIXL1 (Mesendodermal lineage), SOX17 (definitive endoderm), and NCAM1 (mesoderm). (**c**) Majority of SOX17 cells were also FOXA2^+^. (**d**) Fold change of hPSCs for FOXA2 (n = 3 each; 4 vs. 7, p < 0.001; 7 vs. 10, p < 0.001; 4 vs. 10, p < 0.001), SOX17 (n = 3 each; 4 vs. 7, p < 0.001; 7 vs. 10, p < 0.001; 4 vs. 10, p = 0.9978) and NCAM1 (n = 3 each; 4 vs. 7, p < 0.001; 7 vs. 10, p < 0.001; 4 vs. 10, p < 0.001). Scale bar = 125 μm for 20X images. All data are mean ± SD. *p < 0.05; **p < 0.01; ***p < 0.001.

**Supplementary Fig. 2:**
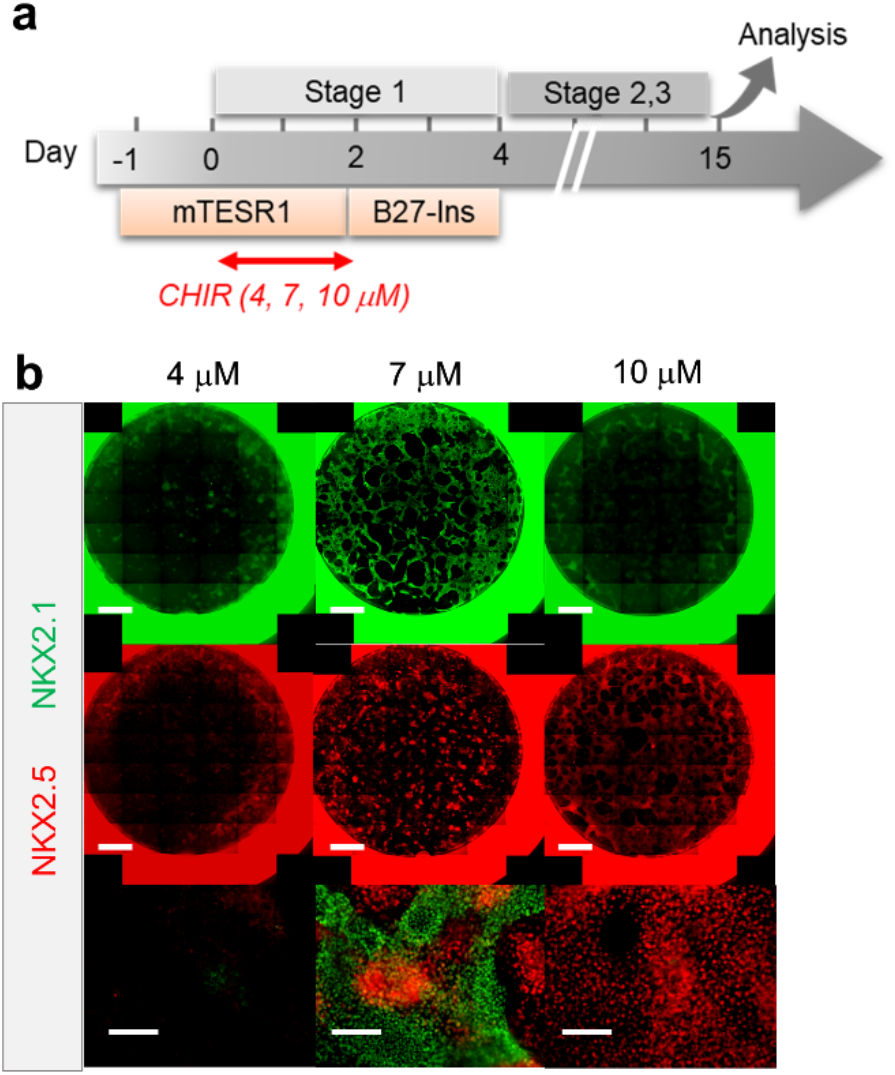
Verification of cardio-pulmonary co-differentiation protocol on BU1 hiPSCs. (**a**) Schematic diagram illustrating the process of cardio-pulmonary co-differentiation, highlighting the adjustment of CHIR concentration during the first 2 days of differentiation. (**b**) IF staining of NKX2.1 and NKX2.5 following 15 days of co-differentiation. Scale bar = 500 μm for whole well scan; Scale bar = 125 μm for 20X images.

**Supplementary Fig. 3:**
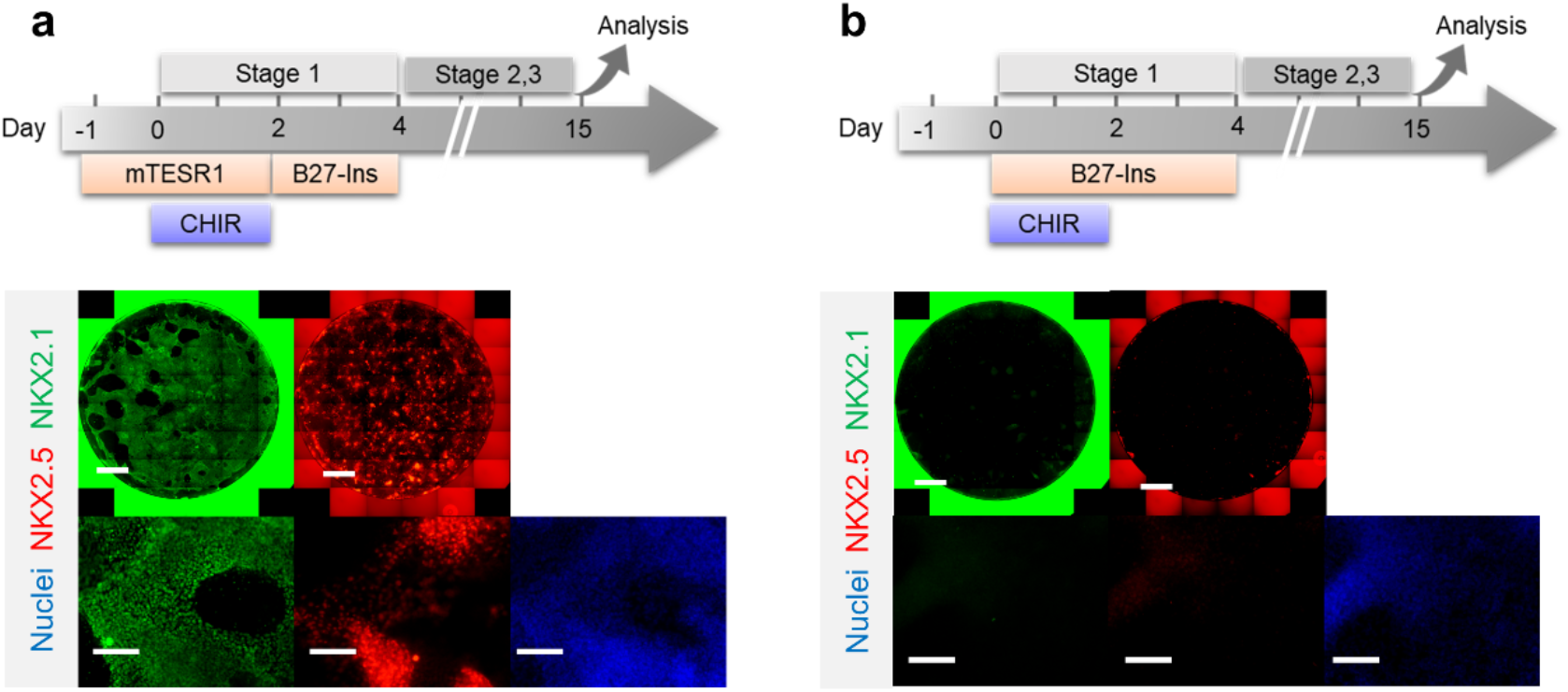
Initial co-induction medium for CHIR-directed differentiation. Cells were induced by CHIR in (**a**) mTESR1 (**b**) and RPMI-based medium, followed by representative IF staining of NKX2.1 and NKX2.5 following 15 days of differentiation. Scale bar = 500 μm for whole well scan; Scale bar = 125 μm for 20X images.

**Supplementary Figure 4:**
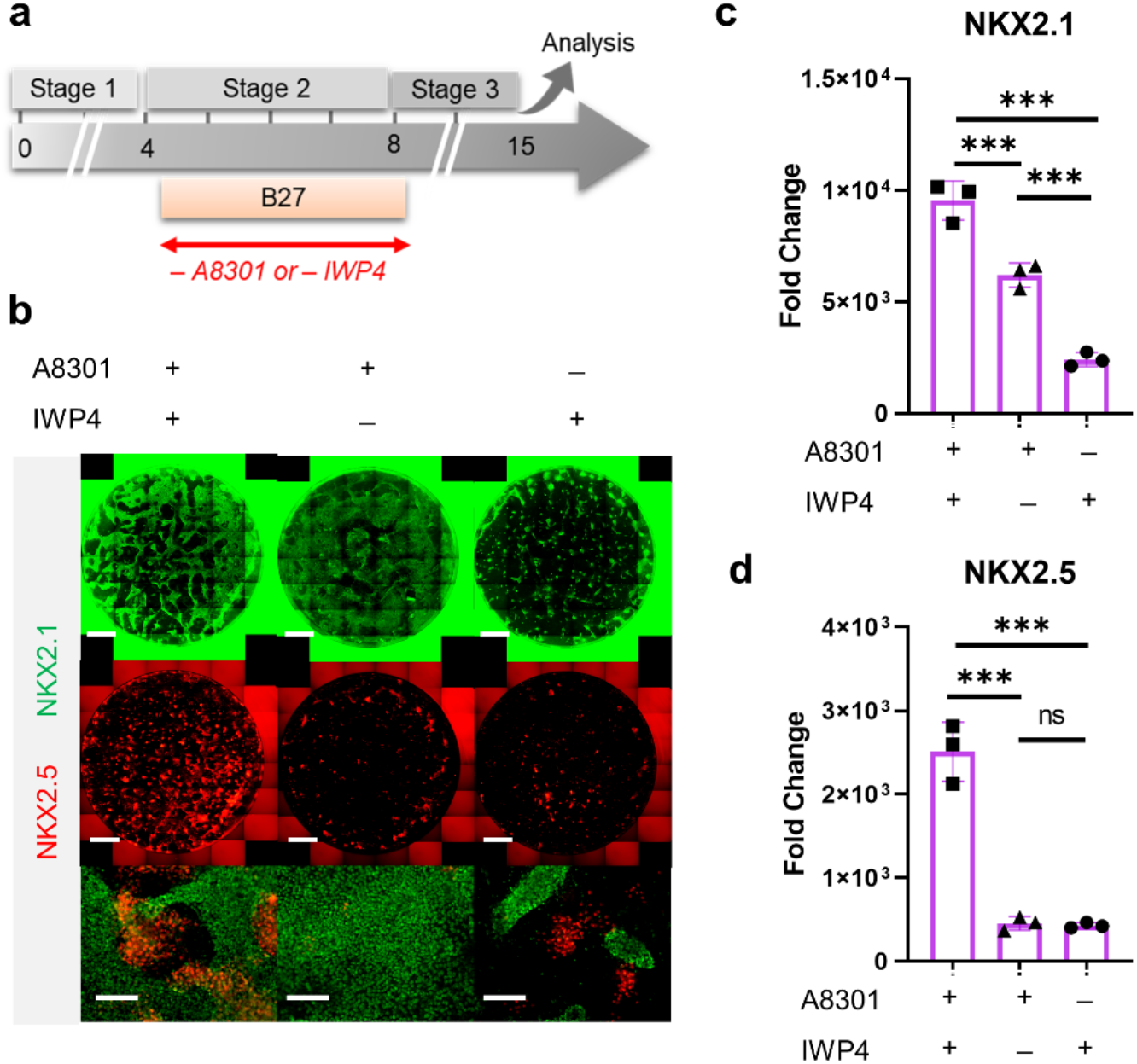
Combination of TGF-β and WNT inhibition during Stage-2 of co-differentiation is required for cardio-pulmonary induction. (**a**) Schematic diagram illustrating the experimental design. (**b-d**) IF staining showing NKX2.1 and NKX2.5 expression on Day 15 of differentiation (**b**), and the corresponding qPCR analysis of (**c**) NKX2.1 (n = 3 each; A8301^+^ /IWP4^+^ vs. A8301^+^ /IWP4^−^, p < 0.001; A8301^+^ /IWP4^+^ vs. A8301^−^ /IWP4^+^, p < 0.001; A8301^+^ /IWP4^−^ vs. A8301^−^ /IWP4^+^, p < 0.001) and (**d**) NKX2.5 (n = 3 each; A8301^+^/IWP4^+^ vs. A8301^+^ /IWP4^−^, p < 0.001; A8301^+^ /IWP4^+^ vs. A8301^−^ /IWP4^+^, p < 0.001; A8301^+^ /IWP4^−^ vs. A8301^−^ /IWP4^+^, p = 0.9986). Scale bar = 500 μm for whole well scan; Scale bar = 125 μm for 20X images. All data are mean ± SD. *p < 0.05; **p < 0.01; ***p < 0.001.

**Supplementary Figure 5:**
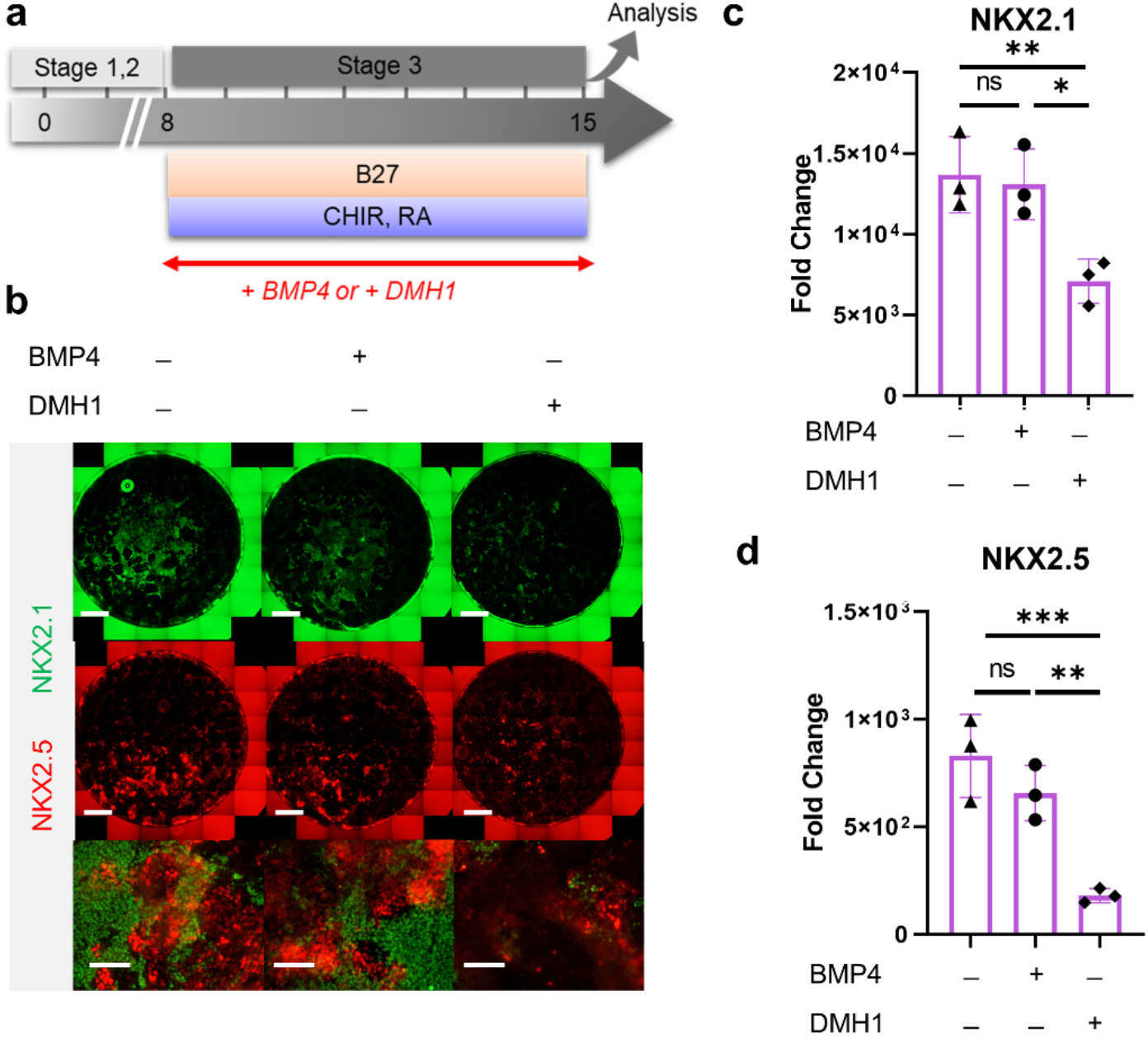
Roles of BMP4 during Stage-3 of co-differentiation. (**a**) Schematic diagram illustrating the experimental design. (**b**) IF staining showing NKX2.1 and NKX2.5 expression on Day 15 of differentiation, and the corresponding qPCR analysis of (**c**) NKX2.1 (n = 3 each; BMP4^−^ /DMH1^−^ vs. BMP4^+^/DMH1^−^, p > 0.05; BMP4^−^ /DMH1^−^ vs. BMP4^−^ /DMH1^+^, p < 0.01; BMP4^+^ /DMH1^−^vs. BMP4^−^ /DMH1^+^, p = 0.9737) and (**d**) NKX2.5 (n = 3 each; BMP4^−^ /DMH1^−^ vs. BMP4^+^ /DMH1^−^, p = 0.3330; BMP4^−^ /DMH1^−^ vs. BMP4^−^ /DMH1^+^, p < 0.001; BMP4^+^ /DMH1^−^vs. BMP4^−^ /DMH1^+^, p < 0.01). Scale bar = 500 μm for whole well scan; Scale bar = 125 μm for 20X images. All data are mean ± SD. *p < 0.05; **p < 0.01; ***p < 0.001.

**Supplementary Fig. 6:**
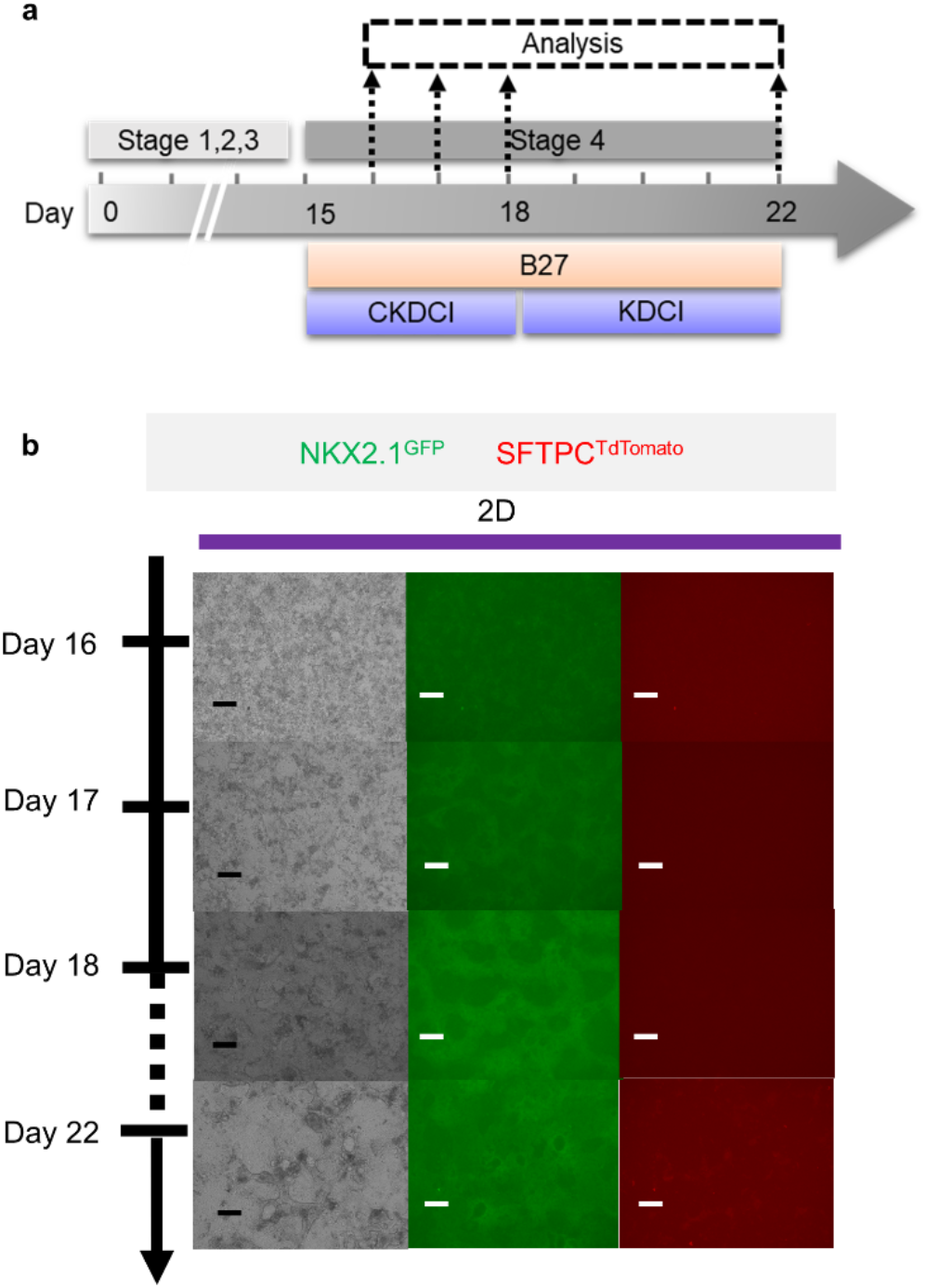
Co-maturation of Day 15 cardiac and pulmonary progenitors on 2D submerged culture. (**a**) Schematic diagram showing the experimental design. (**b**) Live cell imaging of the NKX2.1^GFP^ and SFTPC^TdTomato^ reporter signal over time. Scale bar = 125 μm for 10X images.

**Supplementary Fig. 7:**
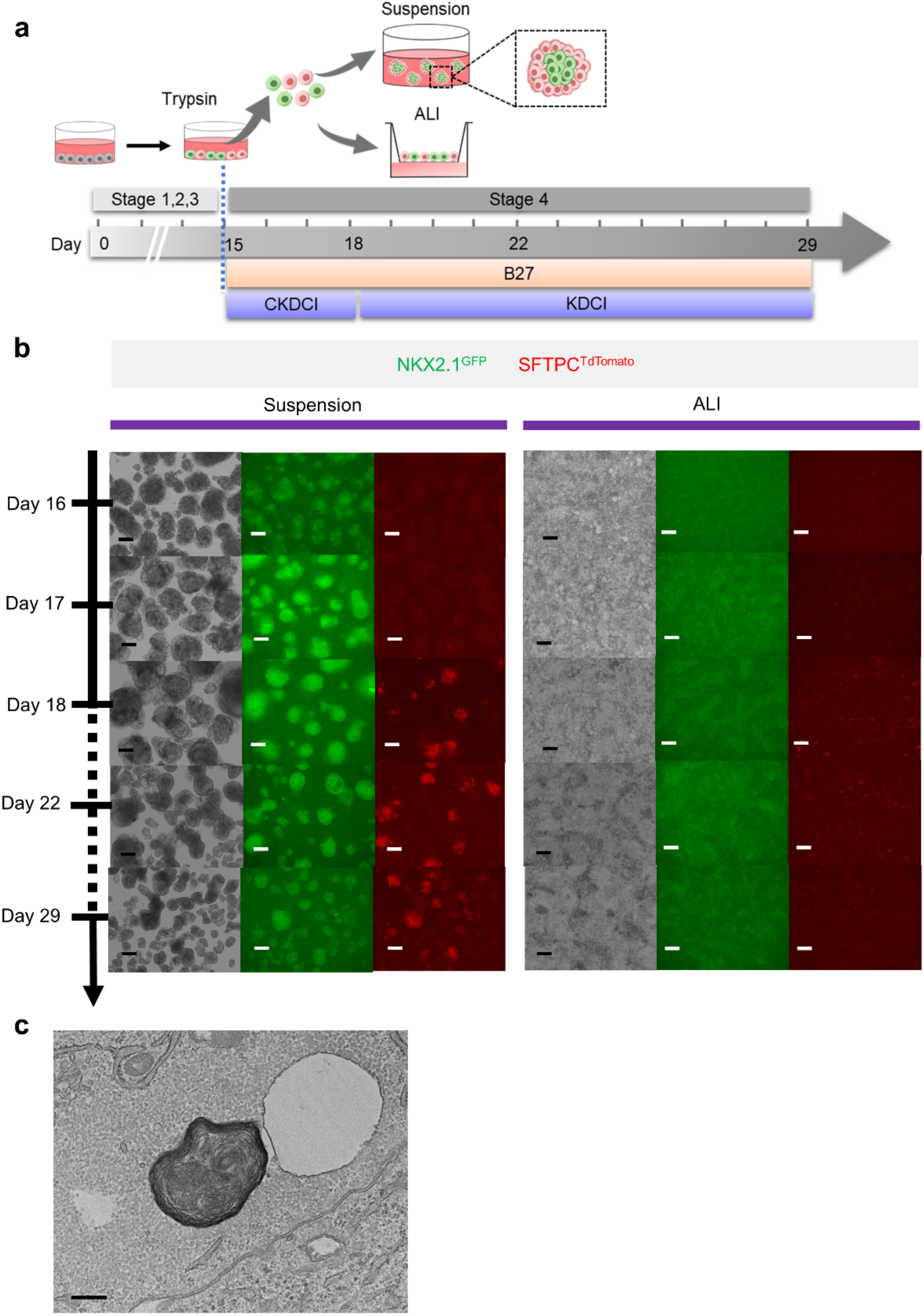
Co-maturation of Day 15 cardiac and pulmonary progenitors on ALI and 3D suspension culture platforms. (**a**) Schematic diagram showing the experimental design. (**b**) Live cell/organoid imaging on the NKX2.1^GFP^ and SFTPC^TdTomato^ reporter signal over time. Scale bar = 125 μm for 10X images. (c) Transmission electron microscopy (TEM) image of the lamellar body of AT2 cells following alveolar maturation in 3D suspension culture, scale: 400 nm.

**Supplementary Fig. 8:**
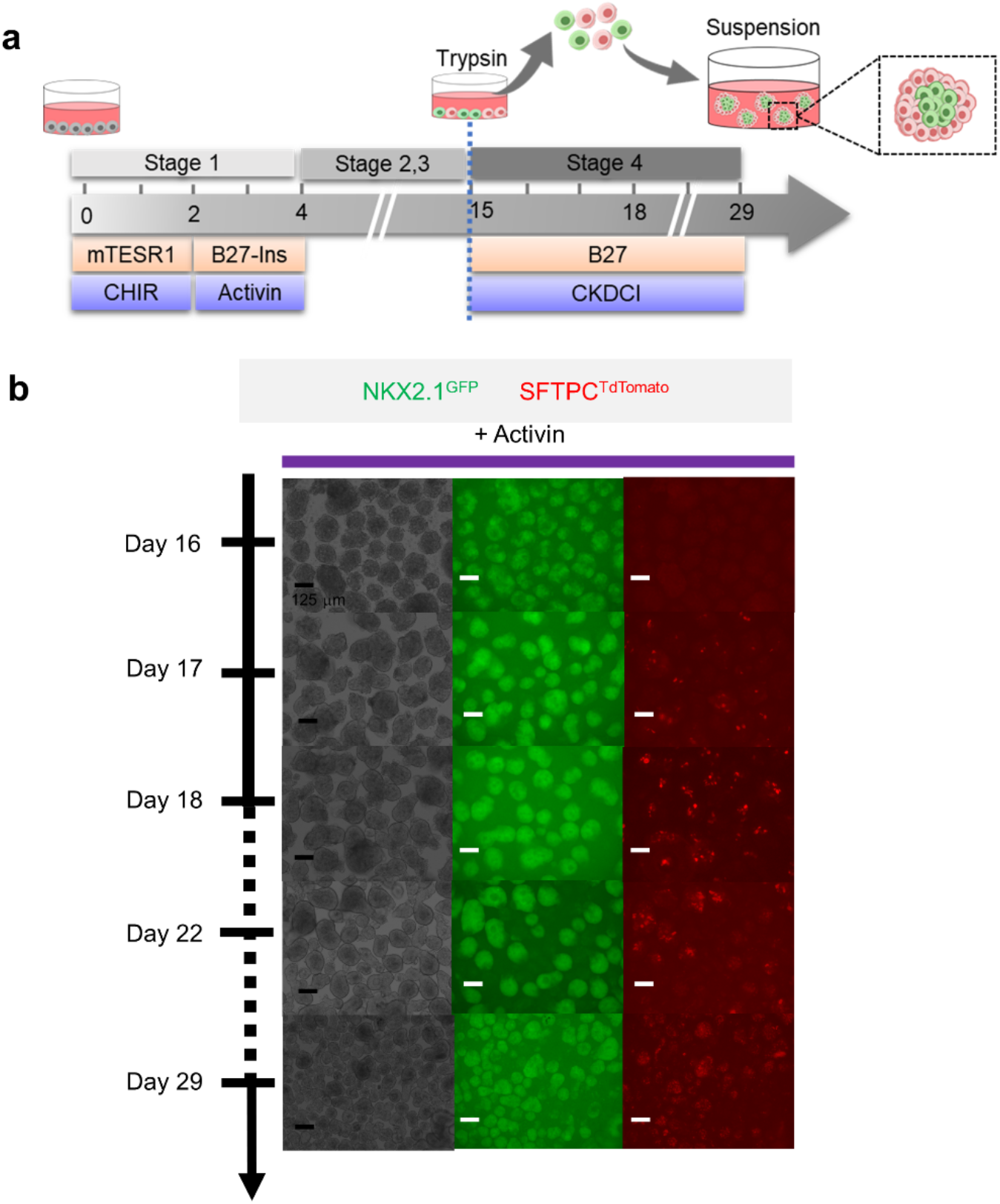
Maturation of pulmonary progenitors derived from Activin A-based protocol on 3D suspension culture. (**a**) Schematic diagram showing the experimental design. (**b**) Live organoid imaging on the NKX2.1^GFP^ and SFTPC^TdTomato^ reporter signal over time. Scale bar = 125 μm for 10X images.

**Supplementary Video 1:** Contacting cardiac μT following 7 days after withdrawal of CHIR.

**Supplementary Video 2:** Calcium influx capability of cardiac μT loaded with Cal-520.

